# A single-cell analysis of the molecular lineage of chordate embryogenesis

**DOI:** 10.1101/2020.03.02.966440

**Authors:** Tengjiao Zhang, Yichi Xu, Kaoru Imai, Teng Fei, Guilin Wang, Bo Dong, Tianwei Yu, Yutaka Satou, Weiyang Shi, Zhirong Bao

**Affiliations:** Institute for Regenerative Medicine, Shanghai East Hospital, School of Life Sciences and Technology, Tongji University, Shanghai 200123, P.R. China; Developmental Biology Program, Sloan Kettering Institute, New York, NY 10065, USA; Department of Biological Sciences, Graduate School of Science, Osaka University, Toyonaka 560-0043, Japan; Department of Biostatistics and Bioinformatics, Emory University, Atlanta, GA 30322, USA; Department of Zoology, Graduate School of Science, Kyoto University, Kyoto 606-8502, Japan; Ministry of Education Key Laboratory of Marine Genetics and Breeding, College of Marine Life Sciences, Ocean University of China, Qingdao 266003, China

## Abstract

In multicellular organisms, a single zygote develops along divergent lineages to produce distinct cell types. What governs these processes is central to the understanding of cell fate specification and stem cell engineering. Here we used the protochordate model *Ciona savignyi* to determine gene expression profiles of every cell of single embryos from fertilization through the onset of gastrulation and provided a comprehensive map of chordate early embryonic lineage specification. We identified 47 cell types across 8 developmental stages up to the 110-cell stage in wild type embryos and 8 fate transformations at the 64-cell stage upon FGF-MAPK inhibition. The identities of all cell types were evidenced by *in situ* expression pattern of marker genes and expected number of cells based on the invariant lineage. We found that, for the majority of asymmetrical cell divisions, the bipotent mother cell shows predominantly the gene signature of one of the daughter fates, with the other daughter being induced by subsequent signaling. Our data further indicated that the asymmetric segregation of mitochondria in some of these divisions does not depend on the concurrent fate inducing FGF-MAPK signaling. In the notochord, which is an evolutionary novelty of chordates, the convergence of cell fate from two disparate lineages revealed modular structure in the gene regulatory network beyond the known master regulator T/*Brachyury*. Comparison to single cell transcriptomes of the early mouse embryo showed a clear match of cell types at the tissue level and supported the hypothesis of developmental-genetic toolkit. This study provides a high-resolution single cell dataset to understand chordate embryogenesis and the relationship between fate trajectories and the cell lineage.

**Highlights:**

- Transcriptome profiles of 47 cell types across 8 stages in early chordate embryo
- Bipotent mother in asymmetric division shows the default daughter fate
- Modular structure of the notochord GRN beyond the known function of T
- Invariant lineage and manual cell isolation provide truth to trajectory analysis

## Introduction

Metazoans possess vastly divergent cell types that develop from a single precursor. Recently, droplet-based high throughput scRNA-seq techniques have been applied extensively to a variety of model systems to study early embryogenesis (Briggs et al., 2018; Cao et al., 2019b; Farrell et al., 2018; Pijuan-Sala et al., 2019; Wagner et al., 2018). These studies inferred developmental paths through trajectory analysis and greatly improved our understanding of how cells differentiate. However, because in these animals the exact cell lineages are not known, the trajectories remain computational hypotheses. Cell barcoding techniques such as those based on CRISPR enable cell lineage tracing (Alemany et al., 2018; Raj et al., 2018) but still face technical limitation in temporal resolution to capture every cell division.

In this regard, model organisms with invariant cell lineage, such as the Nematode *C. elegans* (Packer et al., 2019*) and the protochordate ascidian Ciona*, provide a unique opportunity where the known lineage underlie the interpretation of developmental trajectories. In particular, as the ascidians are sister group to vertebrates, they possess similar body plan and cell types and are crucial to understand how vertebrate developmental programs arose during evolution (Satoh et al., 2003). Recently, a high throughput scRNA-seq study examined *Ciona intestinalis* development from 110-cell to the larva stage, revealing the developmental trajectories of main embryonic cell types (Cao et al., 2019a). However, by the 110-cell stage, the main tissue subtypes, e.g., muscle, heart, etc., have been specified (Figure S1), leaving the questions open regarding the trajectories from the totipotent zygote to tissue specification.

Here we use scRNA-seq to systematically examine lineage specification in early chordate embryogenesis using the *Ciona savignyi* model. Employing manual cell dissociation and isolation, we obtained a total of 750 single cell expression profiles which corresponds to 47 cell types for wild-type and 10 cell types for MEK inhibitor treated embryos. By carefully examining every cell from single embryos, we showed that the number of cells identified in each cell type agrees with known lineage information. These cell type assignments are consistent with *in situ* based cell type classification. We observed prevalent precursor-progeny relationships where the mother cell only possesses the gene signature of one progeny. In particular, we studied the asymmetrical segregation of mitochondria genes and found their asymmetry is unaffected by FGF-MAPK inhibition. Next, we examined the lineage differentiation of notochord and found that two independent lineages contribute to notochord cells through separate gene regulatory cascades that converge on the activation of *Brachyury*. Finally, we performed cross-species comparison of *Ciona* embryonic cell types with those of mouse early embryos and showed that conservation is restricted to a few well characterized master regulator genes, consistent with predictions from previous evo-devo models.

## Results

### Cell isolation and sequencing

To best exploit the invariant cell lineage and low cell numbers in *Ciona* embryos, we dissociated *C. savignyi* embryos at eight developmental stages (1, 2, 4, 8, 16, 32, 64 and 110-cell) and manually collected individual cells from each embryo (Figure 1a). For each embryonic stage, we sampled two to eight embryos for a total of 29 wild-type embryos. Furthermore, we collected cells from two 64-cell stage embryos that were treated with U0126, a MEK inhibitor. We recovered 100% cells from each embryo up to 32-cell stage and more than 90% of cells for 64- and 110-cell embryos, totaling 648 wild-type and 125 U0126-treated cells to be sequenced (Figure 1a,b and Table S1).

**Figure 1.**
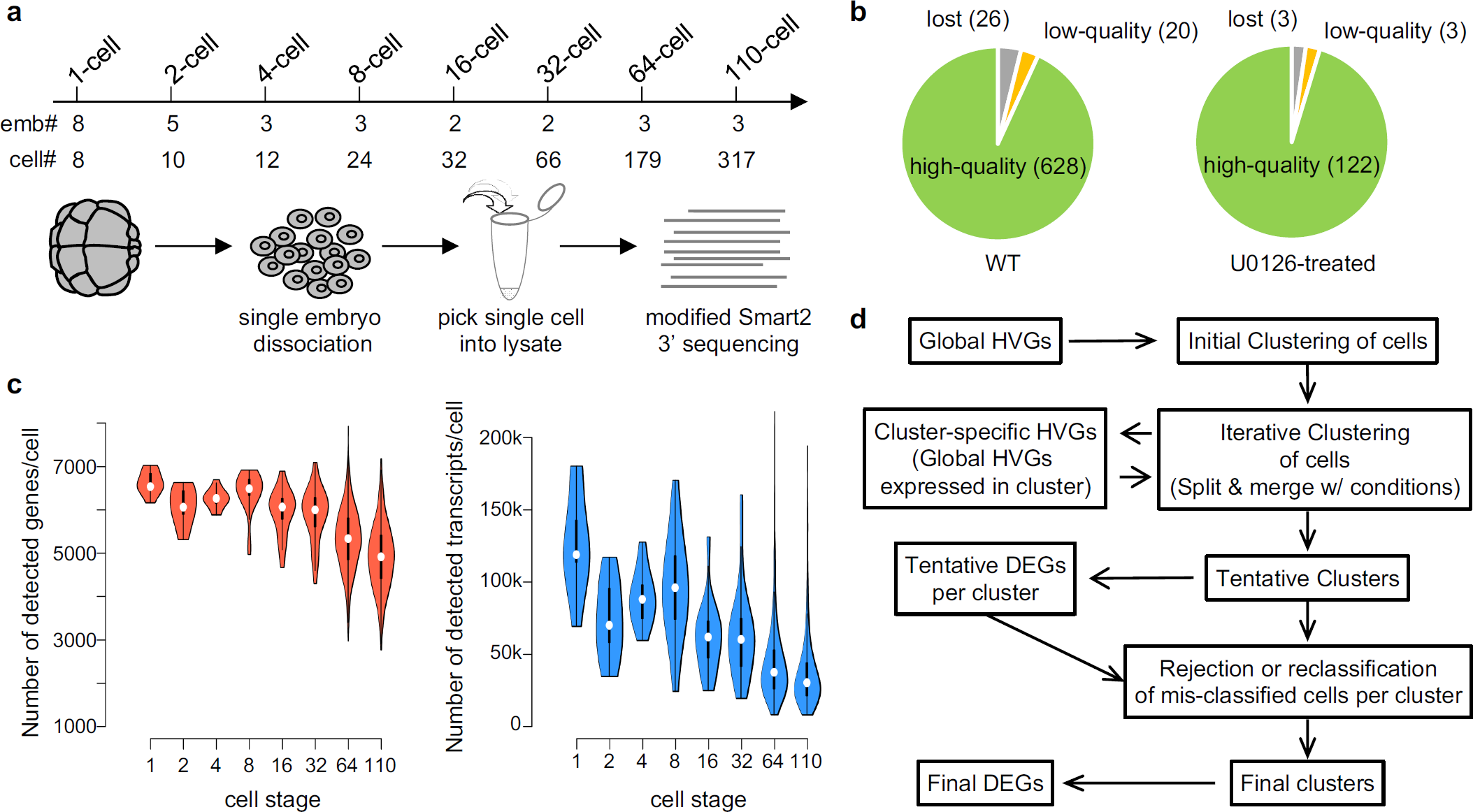
Overview of single-cell RNA-seq assay and cell type classification. a) Number of embryos processed, cells collected and sequenced from the 1- to 110-cell stage. b) Number of cells lost during manual cell picking, showing low-quality transcriptome (<= 2,000 genes), and showing high-quality transcriptome for wild-type (left panel) and U0126-treated (right panel) embryos. c) Distribution of gene (left panel) and transcript (right panel) number per cell at for samples at different developmental stages. d) Computational pipeline for iterative clustering to identify cell types and DEGs.

For each isolated cell, we generated single cell transcriptome using a modified Smart2 protocol that sequences transcripts at the 3’ end and allows transcript counting and multiplexed library preparation. On average, we detected >5,000 genes and a median of 48,340 transcripts per cell (Figure 1c). Earlier stage cells, which have larger cell size, have more transcripts and genes than later stage cells. A small number of cells have less than 2,000 genes detected, which we discarded as low-quality cells. In total, we obtained 750 high quality single cell transcriptomes, including 628 from wild-type and 122 from U0126 treated embryos.

### Cell type identification

For objective identification of cell types, we undertook an iterative clustering approach (Figure 1d) and clustered cells at each developmental stage (Figure S3 and S4). Following identification of highly variable genes (HVGs) from the scRNA-seq data set (see Methods and Figure S2), we used the DBSCAN algorithm (Ester et al., 1996) for unsupervised clustering of cells (see Methods), which not only provides objective measures to optimize for but also allows exclusion of individual cells as outliers (potentially low quality cells). Each cluster undergoes the next round of clustering based on cluster-specific HVGs (Figure 1d). When a cluster is split, we used bootstrapping to examine if the newly produced, tentative clusters are significantly different from each other. Specifically, we measured the similarity of gene expression between two tentative clusters by the accumulating difference of HVG expression (see Methods). If *p* > 0.01, we considered the two groups of cells sufficiently similar and therefore should not be divided. In practice, the majority of the clusters produced showed p-value <= 0.001 (Figure 2b and S3). Because the p value tends to lose significance on clusters with a small number of cells, we accepted some of the smaller clusters with *p* > 0.01 (Figure 2b) after examining the number and quality of different genes expressed in them. The weakest case is the separation of the A-line and B-line notochord cells at the 110-cell stage (p=0.036), which is discussed in details below.

**Figure 2.**
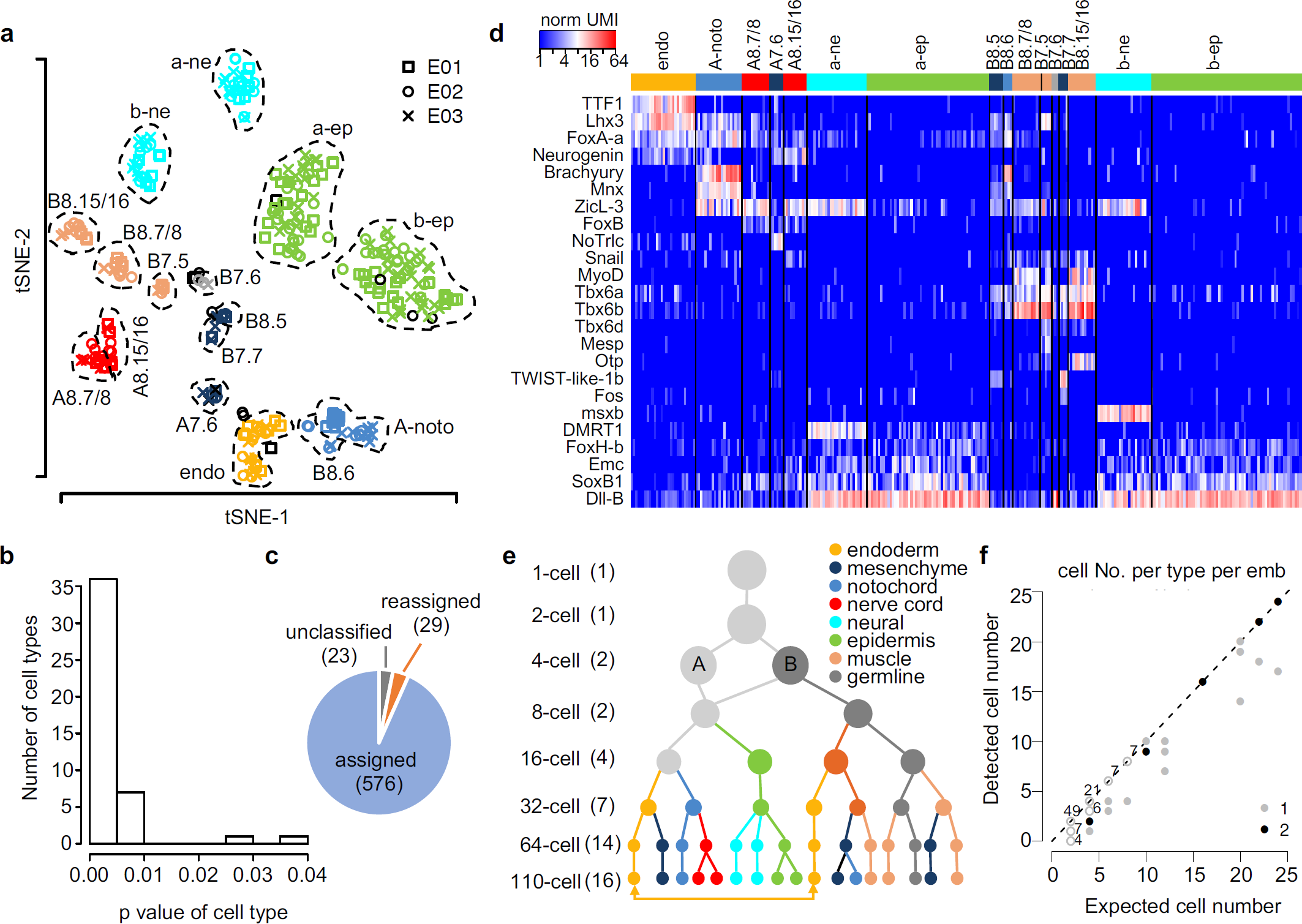
Cell type identification and cell identity assignment. a) Display of identified cell types for the 110-cell stage. Cells from three embryos (E01, E02 and E03) are represented by different symbols (upper right). Clusters are labeled with canonical blastomere names except for a-ne: a-line neural; a-ep: a-line epidermis; b-ne: b-line neural; b-ep: b-line epidermis; endo: A-line and B-line endoderm; A-noto: A-line notochord. Cells of the same cluster are circled by dashed line and colored the same. Black symbols represent rejected/unclassified cells from corresponding embryos. b) Distribution of the statistical significance of the identified cell types across all developmental stages. Statistical significance of a cell type is defined as the p-value of a cell type being the same as the type most similar to it (see Methods). c) Number of cells that are assigned to its original cluster, re-assigned to another cluster or rejected from all clusters across all developmental stages. d) Heatmap of gene expression level for known cell type-specific markers at the 110-cell stage. Each row is a known marker gene. Each column is a cell. Black lines mark the boundaries of cell clusters. Colored bars and names at the top correspond to the clusters in (a). Expression level is shown as Log2 of normalized UMIs. e) Summary of identified cell types in the form of a landscape of differentiation trajectories. Circles represent identified cell types. Lines link progenitors and descendants across adjacent stages. A, B: A and B cell. Double-headed arrow indicates the two nodes of endoderm at 110-cell stage are one indistinguishable type/state. Cell types (color legend, upper right) within the same stage are generally ordered as in the cell lineage. Numbers in parentheses on the left show the number of cell types identified at the corresponding stage. f) The number of cells in each embryo assigned to each type compared to the expected numbers across all embryos and stages. Numbers indicate the number of dots falling at the same coordinates.

A total of 47 cell clusters were defined. These include 2, 2, 4, 7, 14 and 16 clusters for the 4-, 8-, 16-, 32-, 64- and 110-cell stage (Figure 2a and S4), respectively, based on the iterative clustering analysis, while the 1- and 2-cell stage were each accepted as one cluster.

To examine if the assignment of each cell to the corresponding cluster is proper, we performed additional verification by comparing the similarity between the gene expression of a cell and the average expression profile of its tentative cell cluster. We first computed differentially expressed genes (DEGs) for each cluster (see Methods). We then compared gene expression of a cell to the DEGs of its cluster by Pearson’s correlation test (see Methods). A cell is accepted into a cluster if *p* < 10^−5^. A total of 52 cells were reassigned, including 23 that were rejected from all clusters as unclassified cells due to ambiguous gene expression (Figure 2c).

We then determined the blastomere identity of each cluster by examining the expression of known markers of different cell types (Figure 2d) from the closely related species *Ciona intestinalis* (Imai et al., 2004; Imai et al., 2006), and generated the average expression profile for each cell cluster (Table S4) for ensuing expression analysis.

Finally, we ordered the 47 cell types based on their assigned lineage identity across development stages (Figure 2e), which depicts the resolved landscape of lineage differentiation. Notably, our result achieves single-cell resolution of the entire B-line lineage.

As a test for the accuracy of cell type calling, we asked if for each cell type, the number of cells from each embryo agrees with the expected number from the known cell lineage (Figure 2f and Table S2). In all 124 groups of cells from each type and each embryo, the numbers of cells are equal to or less than the expected numbers. For 92 (74%) out of the 124 groups, the number of cells matches exactly the expected number. 33 (97%) out of 34 groups at the 32-cell stage or earlier showed exact match. Most groups with the number of cells less than expected are from the 64- or 110-cell stage where cells were lost during isolation. Considering the fact that the number of cells in a given cell type was not part of the objectives in our computational analysis, the systematic agreement in cell number demonstrates the quality of our cell cluster identification and cell type assignment.

Despite the high success rate in cell type identification, some blastomeres that are known to be distinguishable in *C. intestinalis* are not separated by our iterative clustering. These are limited to two situations where the differences are known to be subtle with only a handful of markers by *in situ* assays, namely early blastomeres at the 8- and 16-cell stages, and tissue subtypes in later embryos (e.g., 110-cell stage a, b-line neuronal subtypes). Several technical issues with our scRNA-seq assay likely contribute to the lack of resolution in these cases. First, some known *C. intestinalis* markers do not have homologs in the *C. savignyi* gene annotation and thus were left out from our sequencing data, such as the early 8- and 16-cell stage marker *Ci-Bz1* used to distinguish a-line from b-line cells (Tokuhiro et al., 2017). Second, some markers are not differentially expressed between known cell types in our data set. For example, *Neurogenin* is expressed in both A8.15 and A8.16 in our data whereas *in situ* only detected expression in A8.16. Table S3 lists all the cases where known cell types were not separated.

We also compared our results with a recently published high throughput single-cell analysis of *C. intestinalis* (Cao et al., 2019a). The published study used the 10x Genomics platform to sample the 110-cell stage at 26x coverage (equivalent of 26 embryos) and reported 14 cell types, compared to 16 cell types in our study from 3 embryos. Specifically, we resolved three tissue subtypes including the A-line nerve cord, B-line mesenchyme and B-line muscle. The separation of these cell types is supported by clear differences in the expression of 11∼31 DEGs and 1∼2 known markers (Figure S5a-c). Meanwhile, our clustering did not resolve the A- and B-line endoderm. The high throughput study revealed six genes that are differentially expressed between the two types (Figure S5d). In our dataset, two of these genes, *EphrinA-a* and *NoTrlc*, were detected robustly in the endoderm cells. Based on the expression level of these two genes, the endoderm cells in our results can be divided manually into a putative A-line group and a putative B-line group. Interestingly, each group has the right number of cells from each embryo. Thus, it appears that the difference reported by the 10x study was marginally detected in our dataset but was not enough to resolve the two cell types by the same statistical threshold for other clusters. Overall, our study achieved comparable power of resolution to the high throughput approach with a 9x higher coverage.

### Identification of DEGs

After identifying the cell types, we characterized the DEGs among them. We took a conservative approach in defining the DEGs by requiring relatively stringent cutoffs for expression level (mean UMI in a cell type > 0.5, fraction of positive cells >= 0.5), fold difference (> 1.8) between expressing and non-expressing cell types, and p-value of Wilcoxon rank-sum test (p < 0.01, Figure S6a,b). In total, we identified 306 DEGs across all developmental stages examined (Figure 3a, S7-9 and Table S5), including most of the known markers. As expected, many of the DEGs are transcription factors (TFs) and signaling molecules (56 and 43, respectively). Among the 387 predicted TFs (Charoensawan et al., 2010), 15% were detected as DEGs. Furthermore, the DEG list contains genes involved in rich and yet unexplored cell-cell interactions (adhesion and other membrane proteins) and context-specific cell biology (cytoskeleton and motor proteins, vesicle trafficking) as well as chromatin related pathways.

**Figure 3.**
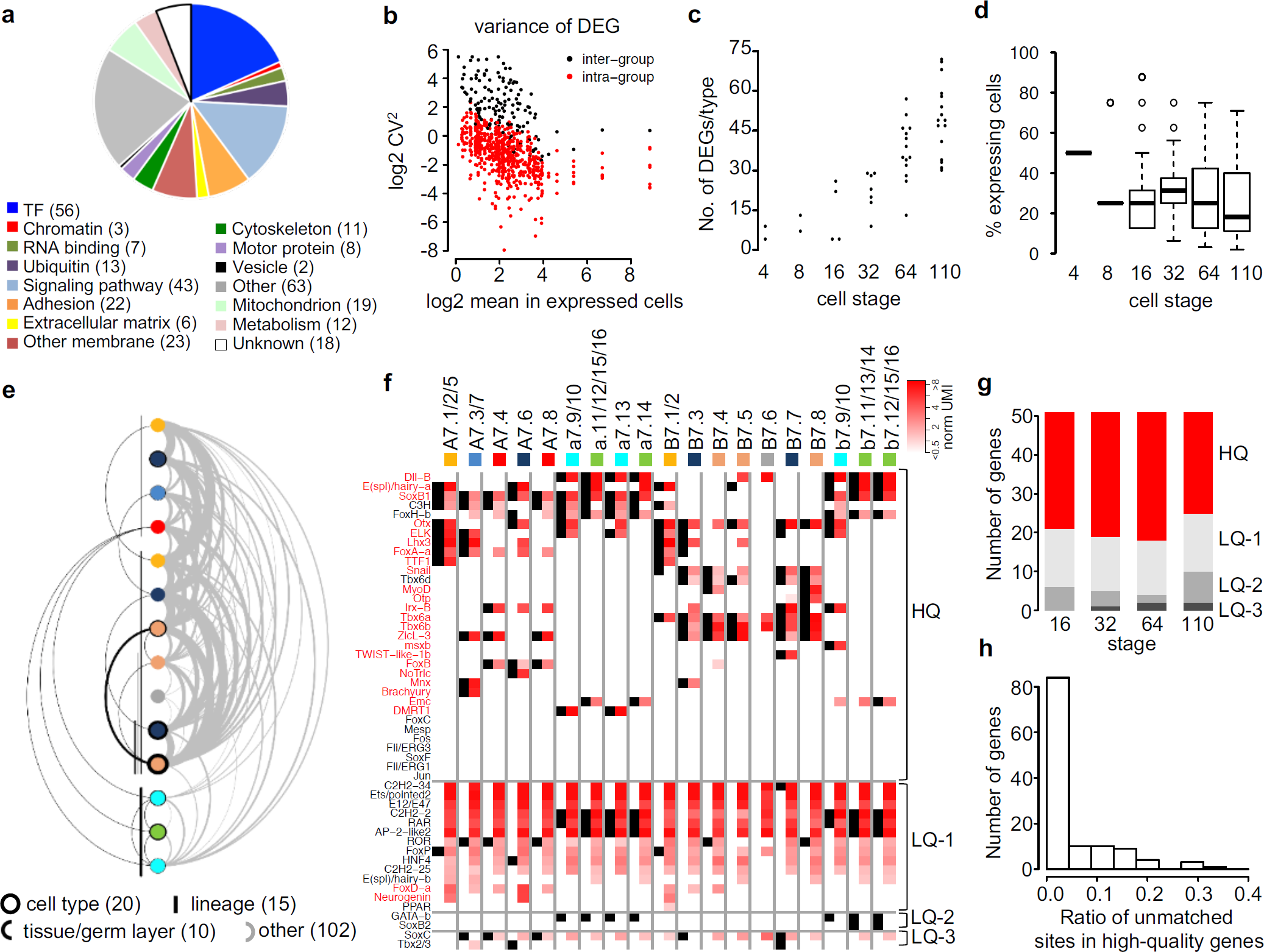
Characterization of DEGs. a) GO term classification of the 306 identified DEGs. b) Variance of expression level among expressing cells (red) and between expressing and low/no expressing cells (black) for the 306 identified DEGs. c) Number of DEGs per cell type across developmental stages. Each dot is a defined cell type. d) Specificity of DEGs as measured by the percentage of expressing cells at each developmental stage. e) Expression pattern of DEGs at the 64-cell stage. Color discs represent cell types, colored as in Figure 2e. From the left, they represent A7.1/2/5, A7.6, A7.3/7, A7.4/8, B7.1/2, B7.3, B7.4, B7.5, B7.6, B7.7, B7.8, a7.9/10/13, a/b epidermis, and b7.9/10, respectively. Black circles around each cell type denote cell type-specific DEGs. Lines underneath denote lineage-specific DEGs. The extent of the lines mark expressing cell types. Arcs link pairs of cells that share DEGs. If a DEG is shared by cells in the same tissue type or germlayer, the arcs are drawn below the cell types. For other DEGs, which have complex combinatorial patterns, are drawn above the cell types. Thickness of lines and circles is proportional to DEG number. DEG numbers in each category in parenthesis. f) Comparison of expression sites for known cell type-specific markers at the 64-cell stage between published *in situ* results (Imai et al., 2006) and detection in this study. Black squares show *in situ* sites. Red squares show detected expression levels by single-cell RNA-seq. Gradient of red (upper right) shows log_2_ of the mean UMI among cells of the corresponding type. HQ: high quality; LQ: low quality. See main text for details. g) Summary of *in situ* and scRNA-seq comparison across embryonic stages. HQ and LQ: see panel f. h) Degree of site discrepancies between *in situ* and scRNA-seq for high quality genes (HQ). Histogram sums genes across embryonic stages.

We then characterized the features of DEGs. The variance of expression level of individual DEGs among the expressing cell types is smaller than and cleanly separated from the variance between the expressing and non-expressing cell types (Figure 3b), demonstrating clear differences in expression levels between the expressing and non-expressing cell types. The number of DEGs per cell type increases over time (Figure 3c), correlating with increased zygotic expression over time and differentiation. In terms of specificity, the DEGs on average are expressed in 25% of the cells at any stage (Figure 3d). Interestingly, the DEGs exhibit complex combinatorial expression patterns. For example, at the 64-cell stage, only 20 out of 147 DEGs are specific to a cell type and 25 are shared by lineage, tissue type or germ layer, while the remaining 102 genes show complex patterns (Figure 3e).

As a comparison of our definition of DEGs, when we applied our DEG-selection thresholds to a mouse E8.25 scRNA-seq dataset (Ibarra-Soria et al., 2018), only 15-33% of the published DEGs passed our thresholds (Figure S6c). Thus, our DEG identification process is more stringent. We reason that a more conservative approach would generate a smaller gene set with clear and reliable patterns, which can in turn serve to nucleate more sensitive searches based on one’s questions and needs.

*In situ* hybridization experiments provide rich and orthogonal information on gene expression patterns. To this end, we compared our results with published *in situ* data of 51 genes in the early *C. intestinalis* embryo (Imai et al., 2006) that have clear homologs in *C. savignyi* (referred to as Imai genes below). We evaluated two aspects of our data: DEG calling and consistency of expressed sites.

First, among the 37 genes that showed differential expression by *in situ*, 23, or 62%, are identified as DEGs in our data (Figure 3f, gene names in red). Among the 14 genes that did not show differential expression by *in situ*, 12, or 86%, are not defined as DEGs in our data. Second, we compared the consistency of expression sites between the two datasets. Taking each gene in each cell type as an expression site, only 8% of expression sites of our defined DEGs show discrepancy with those of *in situ* (Figure 3f, gene names in red). We then focused on 51 Imai genes at the 64-cell stage, among which 33 genes show largely consistent patterns (Figure 3f and S10, HQ/high-quality), including *TTF*-1, *Brachyury, MyoD* in the endoderm, notochord and muscle lineages. Slight differences also exist, such as for *Lhx3*, which is detected by scRNA-seq but not by *in situ* in B7.5, the TVC precursor. Similarly, *Lhx3* is detected in A7.6 by scRNA-seq but not by *in situ*. We observed the opposite pattern as well, such as *E(spl)/hairy-a* being detected by *in situ* but not by scRNA-seq in B7.5. Interestingly, *Lhx3* was reported to be required for TVC specification (Christiaen et al., 2009a), which supports the detection by scRNA-seq. For all stages, about 60% of the Imai genes show high degree of agreement between the two detection methods (Figure 3g). Among these high-quality genes, 78% display <10% unmatched sites between *in situ* and scRNA-seq (Figure 3h). Orthogonal evidence, such as genetic analysis in the case of *Lhx3* in B7.5, is needed to resolve the discrepancy. About 40% of the Imai genes show relatively large inconsistency (Figure 3f), which can be further divided into three situations. At 64-cell stage, 14 Imai genes showed predominantly false-positive sites (Figure 3f, LQ-1) in our data, which show cell-type specificity by *in situ* but were ubiquitously detected by scRNA-seq in almost all cell types. These could be due to errors in the gene models used for *C. savignyi* or erratic RT-PCR. Two Imai genes (*GATA-b* and *SoxB2*, Figure 3f, LQ-2) were not detected by scRNA-seq in any cell type, likely resulted from low sensitivity. Lastly, two Imai genes (*SoxC* and *Tbx2/3*, Figure 3f, LQ-3) showed large fraction of both false positive and false negative detections compared to *in situ* data across different cell types.

Altogether, the systematic comparison between the scRNA-seq and *in situ* data shows moderate sensitivity in DEG detection by scRNA-seq, but high specificity in those detected.

### Insights on differential gene regulation in early ascidian embryo

Initiation of zygotic transcription in the early embryo is a significant step during development. Ascidian embryogenesis does not display global maternal-to-zygotic activation. The earliest detected zygotic expression includes *FoxA-a* and *SoxB1* at the 8-cell stage (Lamy et al., 2006; Miya and Nishida, 2003). We used the scRNA-seq data to systematically examine the earliest zygotic transcription. Specifically, we analyzed the expression level of DEGs between each mother-daughter pair to identify presumptive *de novo* transcription from 1-cell to 16-cell stage (Figure 4a), using the following criteria: a) not detected in mother or ancestors (average UMI in cell type < 0.2 OR total UMI in cell type <=2); b) robust detection in self (average UMI > 0.5 AND fraction of positive cells in type >= 0.5), or present in self (average UMI > 0.25) but robust detection in one of its daughters. We detected extensive *de novo* transcription at the 16-cell stage (Figure 4b and S11), which includes 11 zygotic genes in all three somatic lineages, including the known cases of *SoxB1* and *FGF9/16/20* (Figure S11). However, it is a cell cycle later than the reported initiation at the 8-cell stage. We did detect a putative *de novo* transcription event at the 8-cell stage in the B4.1 cell, which turned out to be a mitochondrial tRNA gene (asterisk in Figure 4b), a likely false positive classification of a maternal gene. The delayed detection in our analysis may result from a combination of our stringent cutoffs and the lack of sensitivity in scRNA-seq.

**Figure 4.**
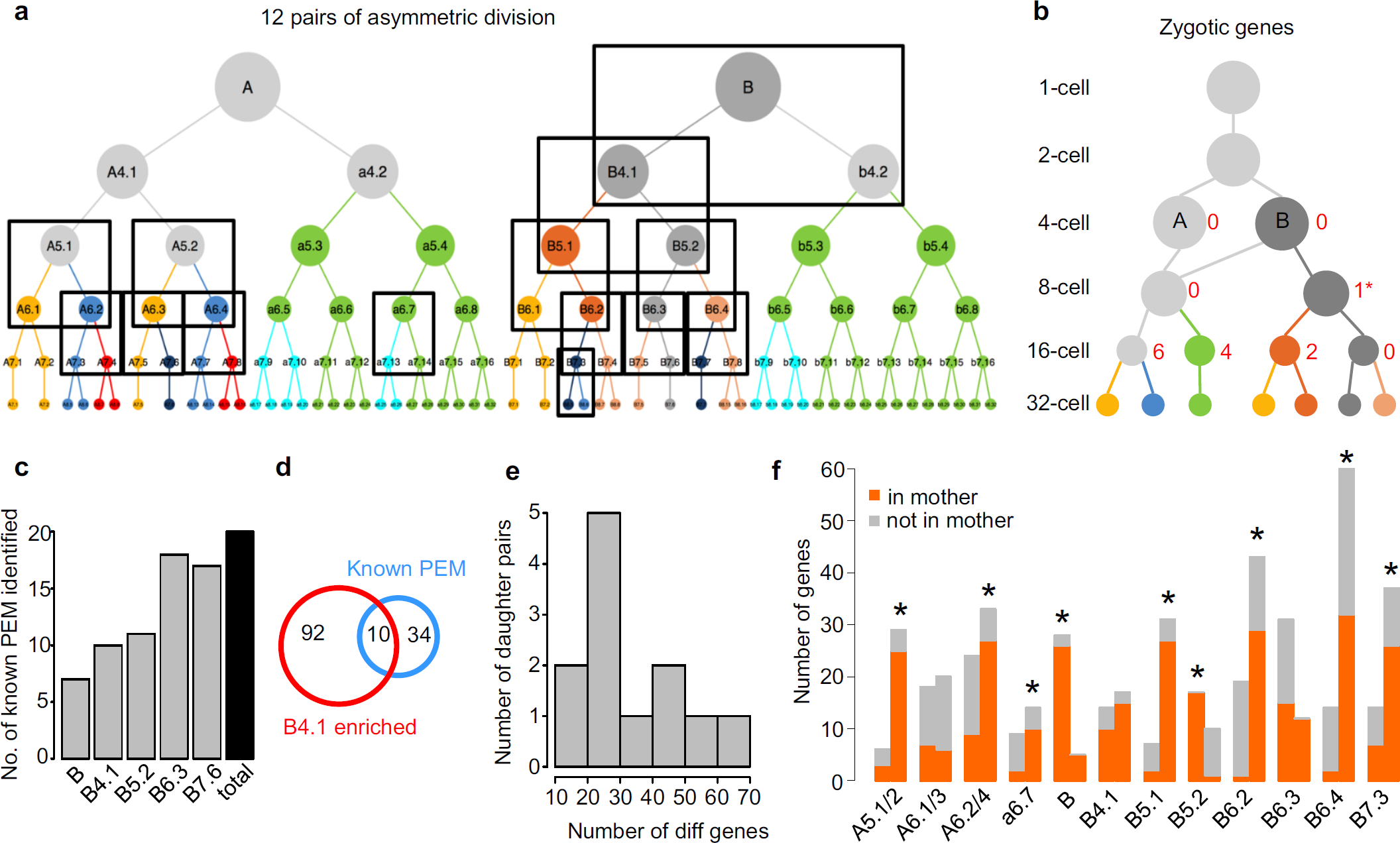
Differential gene regulation in the early embryo. a) 12 asymmetric divisions (boxes, 3 degenerate pairs for A5.1/2, A6.1/3, A6.2/4) resolved by cell type identification. For an enlarged version of the lineage, see Figure S1. b) Number of genes predicted to be zygotically expressed in cells up to the 16-cell stage. Circles, lines and colors follow the scheme in Figure 2e. Numbers next to circles show the number of predicted genes in that cell type. * denotes a false positive prediction (see main text). c) Number of known PEM genes detected by germline enrichment analysis across different stages. d) Overlap between known PEM genes and genes enriched in the B4.1 cell type. e) Histogram of the number of daughter pairs in the 12 asymmetric divisions based on the number of DEGs with >2 fold difference in expression level between each daughter pair. f) Number of DEGs with >2 fold difference in expression level between each daughter pair in each daughter that are detected in the mother cell (orange) or not (grey). Blastomere identities denote the mother cell and each pair of bars represent the two daughters as ordered in the lineage in a.

Before the start of zygotic transcription, the early *Ciona* blastomeres nevertheless display differential gene expression through asymmetric localization/inheritance of maternal mRNA. We detected 26 and 37 DEGs at the 4- and 8-cell stage, respectively (Figure S7). The majority of them show higher number of average UMIs in the germline (B and B4.1), including known cases like *Eph1, Wnt5* and *Cs-pem* (Yamada et al., 2005). However, at each stage there is also a group of genes showing the opposite pattern, such as *FoxJ2* and *in2/Ci-ZF487* with higher level in A and *Ci-ZF087* higher in (the unresolved) A4.1/a4.2/b4.2.

We further examined genes associated with the germline lineage. In ascidian embryo, a group of maternal RNAs called *postplasmic/PEM* are preferentially localized to the posterior blastomeres and contribute to the development of the germline lineage (Prodon et al., 2007). We examined how well the known *postplasmic/PEM* genes can be detected by the pattern of elevated levels in the germline, using a threshold of > 2 fold higher expression level in the germline than its sister lineage across all stages. Of the 44 *C. savignyi* genes that have been annotated as *postplasmic/PEM* (Prodon et al., 2007), 20 can be identified (Figure 4c and S12). In particular, at the 8-cell stage, 10 known *postplasmic/PEM* genes are among the 102 genes enriched in B4.1 (Figure 4d). Previous work showed that some *postplasmic/PEM* genes are both ubiquitously expressed in the cytoplasm of all cells and enriched in a specialized cytoplasmic region (called the CAB) in the posterior cell. Such a pattern may not be reflected as a simple whole-cell elevation in the germline. For the remaing 92 of the 102 genes enriched in B4.1, it is difficult to say if all of these are *postplasmic/PEM* genes without further evidence such as localization to the CAB.

More broadly, we examined asymmetric cell fate specification in general where sister cells become different. In total, we identified 12 types of asymmetric cell divisions where daughter cells were clustered into distinct cell types (Figure 4a, boxes). The daughter pairs showed 10-70 DEGs with >2 fold difference in expression level (Figure 4e). We then asked for each case whether the mother cell, which is in theory bipotent, exhibits characteristic gene expression of the two daughter fates (Figure 4f). The DEGs with >2 fold difference in one daughter compared to the other is considered the characteristic gene expression of the former. We found that the 12 cases fell in two groups. In 3 of the 12, namely A6.1/3, B4.1, B6.3, the mother cell possesses about equal number of characteristic genes of each daughter. In contrast, in the other 9 cases (Figure 4f, asterisk, Chi-squared test p value < 0.01), the mother cell is heavily biased towards one daughter. Among these, two are influenced by maternal determinants of the germline (B, B5.2). The remaining 7 cases except B7.3 are known cases of FGF-induced differentiation. Interestingly, the favored daughters are uniformly the default fate. In the most extreme case of C32 animal cells (a6.7), the mother showed none of the characteristic genes from the induced neural daughter (a7.13). This bias suggests that in the induced daughter the FGF-MAPK signaling would need to suppress the characteristic genes of the default fate, which have already been established in the biased mother. However, not every FGF-dependent induction follows this pattern. In the A6.3 division where FGF-MAPK signaling is required to suppress the A7.6 mesenchymal fate in the A7.5 endoderm, both daughters express comparable number of DEGs inherited from the mother. Thus, FGF-MAPK induction of cell fates may act through different modes of gene regulation.

### The molecular lineage and temporal dynamics of differentiation

Combining the invariant cell lineage and the DEGs in each cell type, we constructed a molecular lineage (Figure 5a), which reveals a global view of how gene expression underlies lineage differentiation, i.e., emergence and turnover of DEGs along lineages. We focused on two interesting cases of lineage differentiation: the divergence of the epidermal lineage and the convergence of the notochord lineages.

**Figure 5.**
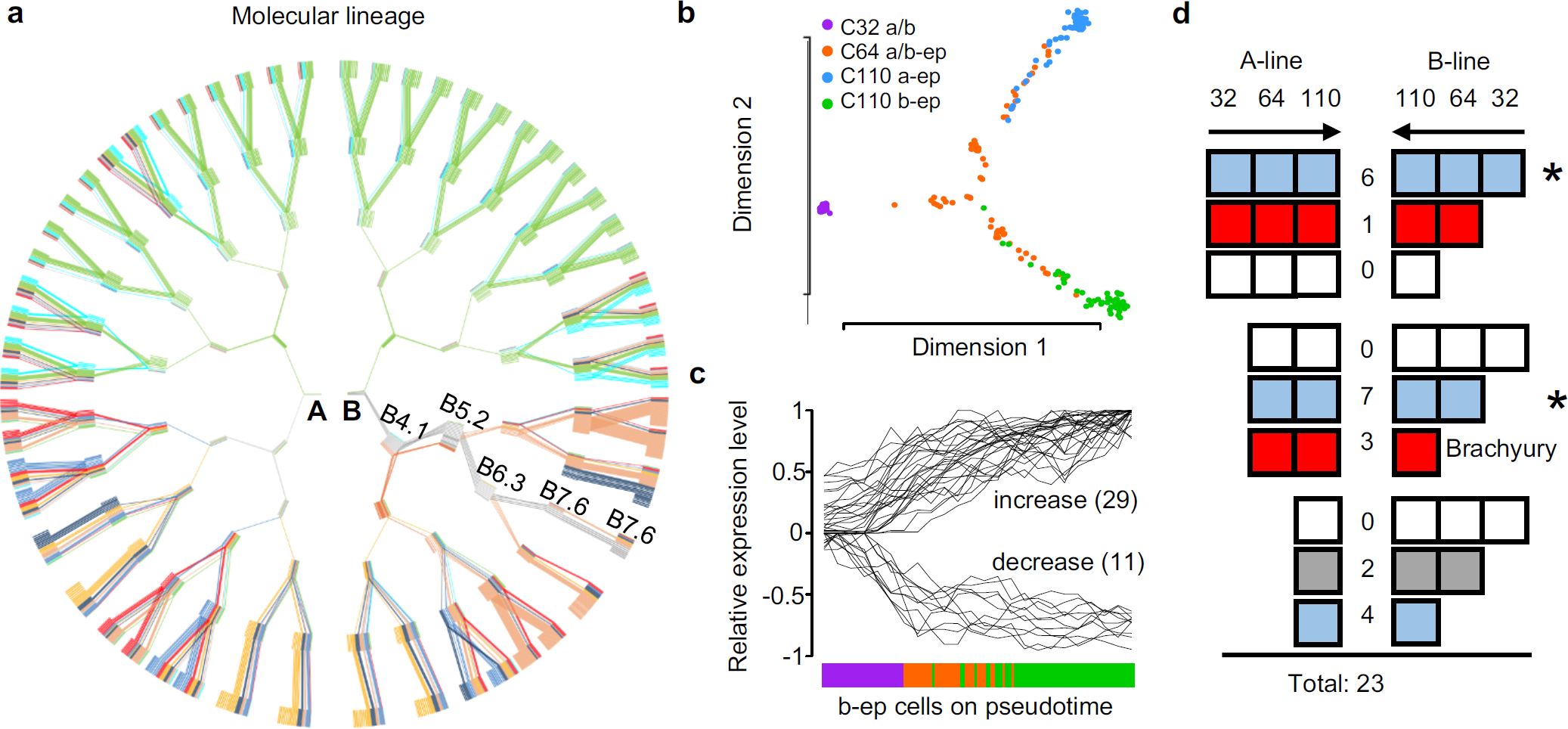
The molecular lineage and temporal dynamics of lineage differentiation. a) Illustration of DEGs in each cell type and their changes across development. Each short colored line in a rectangle node represents a DEG, and each rectangle node (formed by a group short colored lines) represents a blastomere in the cell lineage. Embryonic stages start at the center at 4-cell stage. Identity of the A cell, B cell and cells in the germline lineage are shown as examples. Long lines across stages trace DEGs shared by mother and daughter cells. Colors are based on where a gene is most prominently expressed. Color scheme follows Figure 2e. b) Pseudotime analysis of epidermal fate differentiation at the 32-, 64- and 110-cell stages. Each dot is a cell. C32 a/b: cell type corresponding to the a-line and b-line cells at the 32-cell stage; C64 a/b-ep: cell type corresponding to the a-line and b-line epidermal cells at the 64-cell stage; C110 a-ep: cell type corresponding to the a-line epidermal cells at the 110-cell stage; C110 b-ep: cell type corresponding to the b-line epidermal cells at the 110-cell stage. c) Change of DEG expression level from C32 a/b to C110 b-ep cells (color scheme, see panel b). Cells in C32 a/b, C64 a/b-ep before the bifurcation in panel b, and in C110 b-ep are ordered based on their coordinates on Dimension 1 in panel b to create the x-axis here. d) Comparison of temporal dynamics of notochord-specific DEGs in A-line and B-line. The horizontal range of each rectangle represents the generations of cells (32-, 64- and 110-cell stage). Each pair of rectangles denote the expression pattern of a gene in A-line and B-line (robust expression in which generation). Numbers in between show the number of genes with the corresponding pattern. *Brachyury* expression falls into the pattern colored in red (robust expression starting in A64 and B110). Blue marks the stage-coupled pattern. Grey marks other patterns with 1 or more genes. Asterisks mark permissive modules independent of *Brachyury*.

The a-line and b-line epidermal cells remain as one cell type up to the 64-cell stage but become two distinct types by the 110-cell stage (Figure 2e). To further examine the divergence of cell fates, we used pseudo-time analysis (see Methods) to reconstruct the bifurcation of cellular state toward the a-line and b-line epidermal fates and the accompanying changes of DEG expression (Figure 5b,c). The result showed that the divergence occurs during the 64-cell stage. Furthermore, the analysis showed a leap of cell state from the 32-cell stage to the 64-cell stage, which is followed by gradual transition during the 64- and 110-cell stage (Figure 5b). Consistently, the expression level of DEGs changes rapidly between the 32- and 64-cell stages, but less so between 64- and 110-cell stages (Figure 5c). The leap is not an artifact of smaller number of cells at the 32-cell stage since there is comparable number of cells in each of the gradually changing branches at the 64-cell stage. This pattern is also not an artifact of embryo age variation: cells from individual embryos do not display biased ordering along the pseudo-time (Figure S13a). The abrupt change of cell state indicates rapid gene expression and turnover, which may be dictated by the short cell cycle and continuous differentiation in an invariant cell lineage.

The notochord is considered a chordate invention in evolution (Annona et al., 2015). A key regulator in the gene regulatory network (GRN) is *Brachyury*, which drives notochord differentiation (Satoh et al., 2012). In *Ciona*, the notochord comes from two distinct lineages: the primary notochord from the A-line neural/notochord precursor (A6.2/4) and the secondary notochord from the B-line mesenchyme/notochord progenitor (B7.3). The two notochord lineages become fate restricted at different times: 64-cell stage for the A-line and 110-cell stage for the B-line. The heterochrony raises interesting questions as to if and how the GRN can be launched in different manners.

First, we asked if the temporal dynamics of notochord fate-specific genes are more associated with the embryonic stage or the timeline of notochord fate restriction in each lineage. We identified 23 notochord fate-specific genes, which are defined as genes that are expressed in the notochord lineage and show specificity (>2 fold difference) compared to its non-notochord sister lineage (A7.3/7 vs A7.4/8, A8.5/6/13/14 vs. A8.7/8/15/16, and B8.6 vs B8.5). These notochord fate-specific genes showed three patterns of temporal dynamics (Figure 5d and S13b). The first and predominant pattern is associated with the embryonic stage, where 17 genes are turned on at the same stage in both lineages (Figure 5d, blue). The second pattern is associated with the timeline of fate restriction (Figure 5d, red), where 3 genes are turned on at the notochord fate restriction point (64-cell A-line and 110-cell B-line) and 1 gene in their mothers. Not surprisingly, this pattern identifies the key TFs in regulating the notochord fate, including *Brachyury* and *FoxA-a*. The third pattern is neither associated with embryonic stage nor notochord fate restriction (Figure 5d, grey). These 2 genes are interestingly turned on earlier in the B-line.

Out of the 17 genes associated with embryonic stage, 13 (Figure 5d, asterisk) are expressed in the multipotent progenitors of notochord. Since these genes are turned on before *Brachyury*, they are unlikely to be regulated by *Brachyury*. Furthermore, because these genes are turned on in the multipotent progenitors and show higher expression in the notochord than their sister fates, functionally they are likely permissive factors in establishing the notochord fate. Thus, these results suggest multiple modules in the GRN: the permissive module(s) is/are independent of *Brachyury* and coupled to the embryonic stage; while the timing of fate restriction is achieved by regulating the timing of *Brachyury* expression.

Second, we examined the extent of different gene expression between the A-line and B-line notochord, and asked if the difference reflects their lineage history or diversification of subtypes (Darras and Nishida, 2001; Tanaka et al., 1996). Although A-line and B-line notochord are both restricted to the notochord fate by the 110-cell stage, there are still 21 DEGs that are specific to A-line or B-line (>2 fold difference in A8.5/6/13/14 vs. B8.6). To distinguish between the two models, we examined whether these differences are also present in their progenitors prior to notochord fate restriction. We found most of these differences come from lineage background, with 5 genes including known marker *ZicL* (Yagi et al., 2004) in the A-line vs. 9 genes in the B-line (Figure S13c). In contrast, four genes appear to be newly activated in either the A- or B-line. These genes may contribute to notochord subtype differentiation. Therefore, at this early stage of notochord differentiation, difference in lineage background remains strong between the A-line and B-line notochord but subtypes could be emerging.

### Modes of FGF-MAPK signaling and asymmetric cell division in *Ciona* embryo

Distinct cell types are generated by the interplay between signaling pathways and cellular determinants. The FGF pathway plays broad roles in metazoan development. In *Ciona*, the Fgf9/16/20 dependent MAPK pathway is the major inductive signal for cell fate specification across multiple lineages (Imai et al., 2002a; Kim and Nishida, 2001). We inhibited FGF-MAPK signaling with the MEK inhibitor U0126 (Sakabe et al., 2006) and performed single-cell analysis for two 64-cell embryos (Figure S14 and Table S1).

First, we examined fate transformation upon U0126 treatment. We performed iterative clustering of U0126-treated cells using DEGs from wild-type 64-cell embryo and identified 10 cell types (Figure 6a and S14a,b). Among these, we identified 8 fate transformation events including all 7 known cases such as notochord to nerve cord, and mesenchyme to muscle (Figure S14c). SplitsTree (Huson and Bryant, 2006), which was originally developed for evolutionary phylogeny construction, was used to visualize the similarities of cell types between wild-type and U0126-treated embryos (Figure 6b). Interestingly, we noticed that for the case of notochord and mesenchyme induction, whereas the corresponding FGF-MAPK targets are diminished by U0126 treatment, the presumptive notochord and mesenchyme blastomeres do not adopt a complete fate transformation. They are still separable from the sibling nerve cord and muscle fates by 4 or 5 retained DEGs (Figure S14b), although the differences are much reduced compared to more than 30 DEGs in the WT (Figure 4f). In addition, our analysis revealed a novel fate transformation event in which the presumptive TVC (B7.5) is transformed to a muscle-like fate B7.4/B7.8 (Figure S14c). This is consistent with the previous observation that the expression of *Mesp* in B7.5, a key TF of the TVC fate, is partially dependent on Fgf9/16/20 (Christiaen et al., 2009a). Our result further revealed that upon loss of MAPK signaling B7.5 adopts a muscle fate.

**Figure 6.**
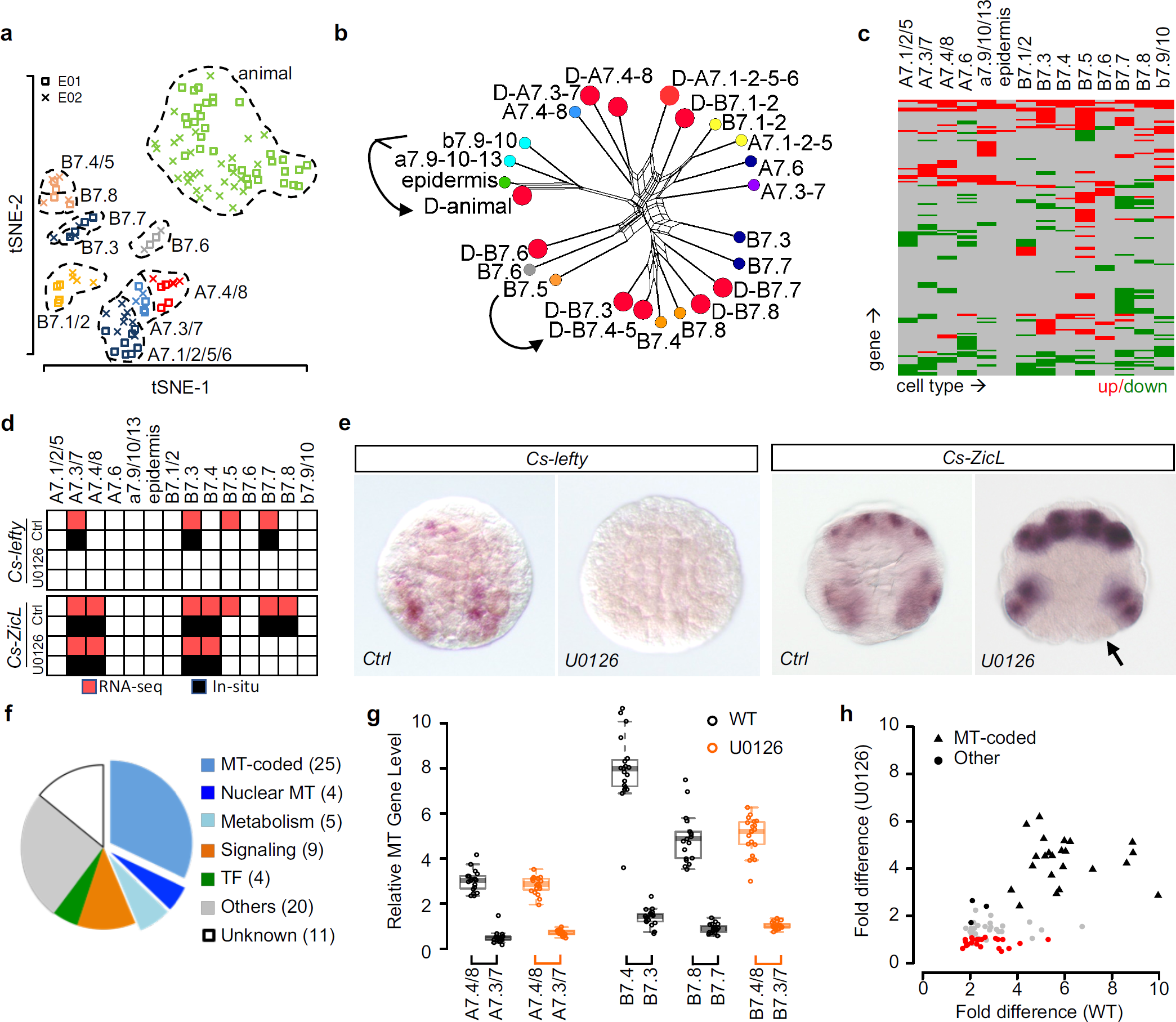
Analysis of FGF-MAPK signaling in fate specification and asymmetric cell divisions. a) Display of 10 identified cell types in 64-cell stage U0126-treated embryos. See Figure 2a for convention. b) SplitsTree showing similarity of DEG profiles among wild-type and U0126 treated cell types. U0126-treated cells are denoted by red dots; wild-type cells colored as in Figure S1. Arrows denote examples of fate transformation. c) Changes of DEG expression at the 64-cell stage after U0126 treatment. See Table S5 for the list of genes and expression levels. d) Summary of detected expression sites at the 64-cell stage by scRNA-seq (red) and *in situ* (black) for *Cs-ZicL* and *Cs-lefty* in wild-type and U0126-treated embryos. e) Representative *in situ* hybridization results of *Cs-lefty* and *Cs-ZicL* in wild-type (Ctrl) and U0126-treated embryos. *Cs-lefty* shows total loss of expression, while *Cs-ZicL* is lost in the posterior most mesodermal lineages (arrow). f) GO term classification of genes co-segregating with mitochondrion (MT)-coded genes in three FGF-dependent sister pairs. g) Asymmetric enrichment of MT-coded genes between sister pairs in wild-type and U0126-treated embryos. h) Comparison of asymmetric enrichment for MT-coded and co-segregating genes between the wild type and U0126 treatment. Each dot is a gene. Black, grey and red: asymmetric in all cell pairs, in some pairs and symmetric after treatment. Fold difference calculated as the geometric mean of fold differences of the relevant cell groups in g.

Next, we examined gene regulation by the FGF-MAPK pathway. U0126 treatment caused broad changes in DEG expression across cell lineages at 22% of the expression sites (Figure 6c and Table S5). Specifically, known FGF-MAPK targets including *Brachyury, Twist-like* and *Otx* were diminished from the corresponding cell types, confirming the specificity of the inhibitor. Our analysis also identified novel FGF-dependent gene expression. For example, *Lefty*, which belongs to the TGF-β superfamily, is normally expressed in notochord, B-line mesenchyme and TVC in WT embryos (Figure 6d). Upon U0126 treatment, the expression is completely diminished in the embryo, suggesting that *Lefty* is a novel target for FGF-MAPK. In addition, we found that *ZicL*, an early specifier for mesoderm lineages, is under FGF-MAPK regulation in a context dependent manner. In U0126-treated embryos, *ZicL* is specifically lost in the posterior muscle and mesenchyme lineage (B7.7/B7.8) but unaffected in the anterior muscle and mesenchyme (B7.3/B7.4) or the A-line blastomeres (Figure 6d). The regulation of *Lefty* and *ZicL* by the FGF-MAPK pathway on was further confirmed by our *in situ* experiments (Figure 6e).

Finally, we examined the interplay between FGF-MAPK signaling and asymmetric inheritance of cytoplasmic determinants during lineage differentiation. Among the asymmetric cell divisions that depend on FGF induction, 3 cases, namely A7.3/7 vs A7.4/8, B7.3 vs B7.4 and B7.7 vs B7.8, are accompanied by transport of mitochondria (MT) towards the marginal daughter (Negishi and Yasuo, 2015), which is reminiscent of asymmetric MT inheritance in mammalian embryonic development and stem cell differentiation (Katayama et al., 2006; Van Blerkom et al., 2000). This pattern can be robustly detected in our scRNA-seq dataset by examining MT-coded genes (Figure S14d). We identified 78 genes during these divisions whose mRNA show enrichment in the marginal daughters compared to their medial sisters. In addition to 25 MT-coded genes, this gene list also contains genes encoding signaling pathways and TFs (Figure 6f). We then asked if the asymmetry of MT and the co-segregating genes requires FGF-MAPK signaling. After U0126 treatment, 24 of the 25 MT-coded genes remain asymmetric (Figure 6g), which suggests that MT segregation does not require FGF-MAPK signaling and another polarity cue exists. Similarly, 31 non-MT genes remain asymmetric. Further experiments are required to determine if these mRNAs are transported by the same mechanism as the MT. Finally, 22 non-MT genes become symmetric (Figure 6h), which are consistent with being conventional target genes of the FGF-MAPK pathway. Taken together, these results suggest that multiple polarity pathways function in these asymmetric cell divisions, and may involve transport of mRNA in addition to MT.

### Systematic comparison of cell types in early chordate embryogenesis

As the sister group to vertebrates, the ascidians provide a critical node in evolution to understand how the vertebrate developmental programs arose. To this end, we conducted a pairwise comparison between *Ciona* and mouse embryonic cell types using our data and a recently published single-cell transcriptomic dataset for the E8.25 mouse embryo (Ibarra-Soria et al., 2018). *Ciona* embryonic tissue specification occurs before gastrulation with the 7 major tissue types emerging by the 110-cell stage, while in vertebrates, such tissue specification events do not happen until after gastrulation (E6.5). Many master transcription factors determining cell fate are expressed commonly in 64-to 110-cell *Ciona* embryos and E6.5 to E8.5 mouse embryos, such as *Nkx2*.*1*/*TTF-1* in the endoderm, *Brachyury* in notochord, *MyoD* in muscle and *Mesp* in cardiac lineages. Thus, the 64- and 110-cell *Ciona* embryo is generally comparable to E6.5 to E8.5 mouse embryo in terms of tissue specification, despite the heterochrony in gastrulation.

To perform cross-species comparison, we first filtered the published mouse DEGs for each cell type with our more stringent thresholds (Figure S6c), reducing the number of DEGs from 3252 to 1788. Globally, we found that comparable fractions of TFs in the *Ciona* and mouse genomes are expressed as DEGs (Figure 7a), and the E8.25 mouse shows a larger fraction of genes in DEGs (Figure 7a). DEGs with orthologs, which are defined by mutual best BLASTP hits (Table S7), are used for further cell type comparison. Surprisingly, only 10% of the mouse DEGs with *Ciona* orthologs have their orthologs detected as DEGs in *Ciona* (Figure 7b). Nevertheless, in terms of gene function, the two sets of DEGs show similar fractions in GO terms that are important for development (Figure 7c). In summary, the two species may activate different genes in the same functional categories for tissue differentiation and morphogenesis.

**Figure 7.**
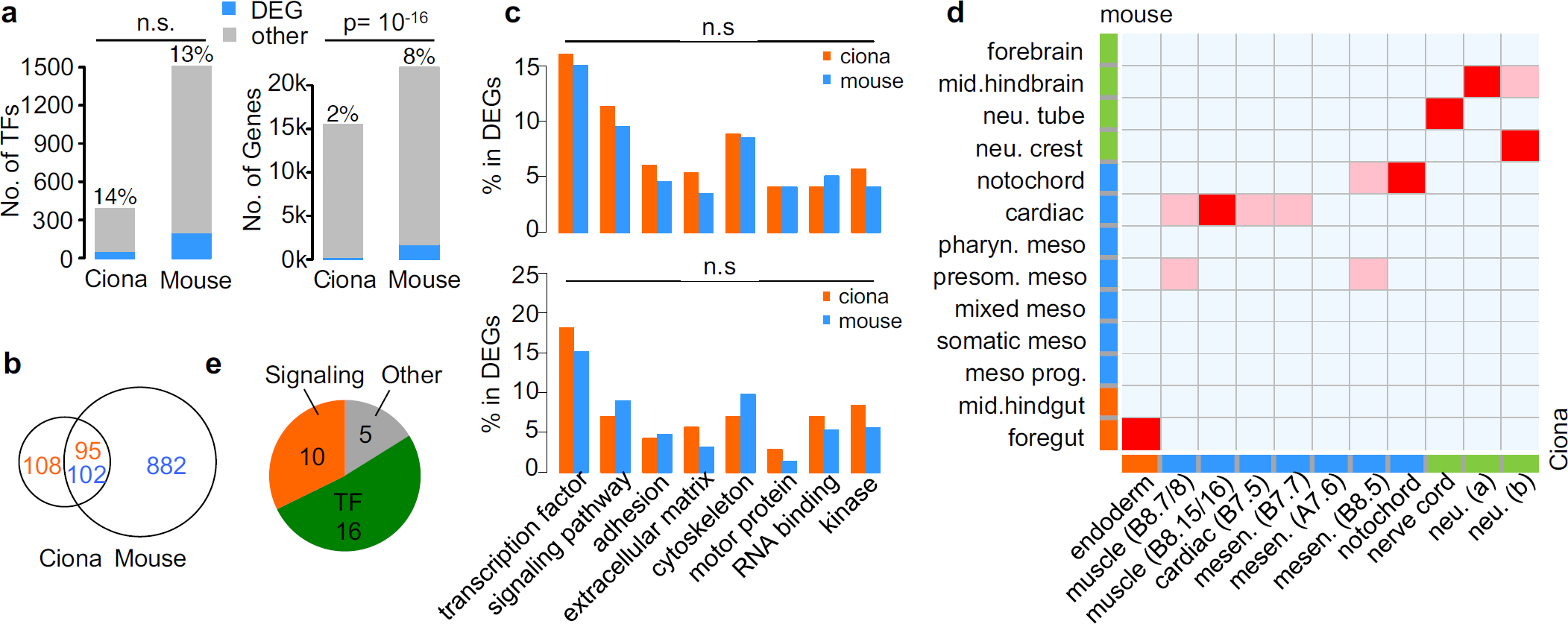
Comparison of single-cell transcriptomes of *Ciona* and mouse embryogenesis. a) Fraction of TFs (left) and all genes (right) that are detected as DEGs in *Ciona* and mouse. b) Number of DEGs with and without orthologs in the other species. c) GO term classification of orthologous DEGs in *Ciona* and mouse. Upper panel: All DEGs with orthologs in the other species; lower panel: Non-overlapping DEGs as in b. d) Matching of cell types based on similarity of DEGs. Red: high similarity; pink: moderate similarity. e) Classification of shared DEGs between matched pairs of cell types in d.

By scoring the type and number of orthologs in DEGs, we were able to match cell types between the 64-/110-cell stage *Ciona* and E8.25 mouse for the majority of the cell lineages (Figure 7d, Table S7, Methods). Six pairs of cell types match with high-similarity and 10 out of 11 *Ciona* cell types have high or moderate similarity to mouse cell types. In all matched pairs, mouse and *Ciona* cell types are derived from the same germ layer. All matched pairs with high-similarity reflect known homologous relationships at sub-tissue level (such as *Ciona* nerve cord and mouse neural tube), except that mouse cardiac tissue has slightly higher similarity to the body wall muscle of *Ciona* than cardiac muscle. Interestingly, the *Ciona* endoderm matches the mouse foregut, but is less similar with hindgut. This is consistent with the previously reported deep conservation of bilaterian foregut gene expression programs (Arendt et al., 2001). Notably, the similarity between cell types is limited to a small number of TFs and signaling molecules (Figure 7e), which is congruent with the hypothesis of developmental-genetic toolkit (Carroll, 2008) and further validates the quality of biological information in scRNA-seq datasets.

## Discussions

Single-cell technologies have revolutionized the developmental biology field and provide an unprecedented opportunity to reveal the role of individual cells in the developing embryo. For most model systems, the lineages are variable, making trajectories as theoretical assumptions. We utilized the invariant lineage of the *Ciona* system to provide a single-cell atlas of early embryogenesis and revealed complex patterns of gene expression and cell fate specification. The expression profiles of *Ciona* embryonic cell types provide valuable resources not only for the study of lineage differentiation but also for the assessment of trajectory reconstruction methods.

Mouth pipette picking was used to prepare single-cell libraries. Each embryo was dissociated separately with cells being carefully checked and counted under microscope before picking, which eliminated the possibility of doublets and consequent false-positive cell types, and the bias of capturing diverse cell types as well. More importantly, accurate cell numbers can be used as the benchmark for cell-type identification, empowering us to study gene dynamics along true differentiation paths defined by the invariant lineage.

One consequence of the manual picking method is a relatively small sample size, limited by the difficulty of capturing every cell of a late-stage embryo and the labor intensity of Smart2 library preparation. However, our *de novo*, unsupervised and objective clustering results turned out to be quite convincing, taking cell numbers of each type as a post-hoc validation. Comprehensive quality controls, including controlling the statistical significance of identified cell types (Figure 2b) and the distance of distribution of DEG expression between expressing and non-expressing cells (Figure 3b), also indicate that our small sample size did not impair the identification of clusters or the definition of DEGs. Compared with the published 110-cell stage dataset with a 9-fold greater sample size (Cao et al., 2019a), our data provides a comparable and perhaps better clustering result (16 vs. 14 cell types). The weakest separation case in our cell types (A-and B-line notochord) is still supported by 21 DEGs including 2 known markers (Figure S13c). For all the known 63 “cell types”, 37 “types” are successfully resolved as separate clusters whereas 26 “types” are collapsed into 10 clusters which are mainly limited to “possible types” of the same biological identity albeit with 1∼3 differential genes (Table S3). Therefore, a relatively small sample size could provide high resolving power.

We systematically evaluated the sensitivity and specificity of our scRNA-seq data. Taking Imai’s *in situ* results as a reference, our scRNA-seq data shows an acceptable level of sensitivity (62%) and a fairly good specificity (86%) at DEG calling (Figure 3f), which means the detection of DEGs is more likely to be true signal rather than noise despite that some DEGs may be missed. Furthermore, the expression patterns of DEGs match well by cell type with those of *in situ*, with only 8% of the sites show discrepancy between two approaches (Figure 3f). It should be pointed out that one of the discrepancies was validated by genetic experiments, where *Lhx3* is detected in the TVC precursor. However, scRNA-seq by design cannot capture all transcripts in the cell especially transcripts in the nuclei. That is part of the reason that our data cannot detect zygotic gene activation at the 8-cell stage, which is still largely nascent transcripts in the nucleus (Figure 4b). Besides, for transcripts with subcellular preference, scRNA-seq shows a poor performance, compared to those methods with cellular structure preserved (Figure 4d). We only observed asymmetrical patterns for a quarter of the *postplasmic/PEM* genes, probably because gene expression levels are averaged out by the whole-cell.

Single-cell analysis allows us to discover new knowledge of expression. Previous genetic study observed that the anterior and posterior endoderm display different properties in experimental perturbation where *NoTrlc*, a previously known specific determinant for A7.6 mesenchyme fate, is expanded into the anterior but not posterior endoderm cells upon FGF-MAPK inhibition (Shi and Levine, 2008). Our analysis, as well as the recently published C. *intestinalis* single-cell study, both detected *NoTrlc* expression at low levels in the anterior endoderm at the 110-cell stage. Thus, it appears that *NoTrlc* is actually expressed in both the endoderm and mesenchyme daughters of the A6.3 precursor cell, but will subsequently be lost in the endoderm lineage while becoming specific to the mesenchyme lineage. This is an example of the classical binary fate choice model whereas both daughters initially express a TF but one cell subsequently loses its expression and the TF becomes highly expressed in its sibling cell.

Asymmetric cell division is a major mechanism of cell fate diversification. By comparing patterns of shared DEGs between the mother-daughter trio, we noticed that in the majority of asymmetric divisions (9/12) the mother cell is heavily biased towards one daughter where one daughter expresses 7-30 fold more mother DEGs than its sibling cells (Figure 4f). Particularly, 6 biased pairs are FGF-MAPK dependent and we found that part of the asymmetrical genes in these divisions became symmetrical upon U0126 treatment, suggesting that FGF-MAPK is required for the patterning of these genes. Further experiments are needed to determine if the mechanism of FGF-MAPK dependent asymmetry is molecular transport in the mother cell or transcriptional repression in the induced daughter. Additionally, there was one group of asymmetrical genes unaffected by U0126, including MT genes and non-MT genes, suggesting a different polarity pathway. Therefore, our results indicate that there are multiple polarity pathways involved in transcriptional regulation and possibly post-transcriptional regulation in asymmetrical cell divisions.

Convergent differentiation, which means one cell type develops from multiple precursor lineages, is seen in the primary and secondary notochord, primary and secondary muscle in *Ciona* embryo development. Interestingly, *Ciona* notochord comes from two distinct lineages that become fate restricted at different embryo stages. The heterochrony allows us to distinguish notochord fate-coupled gene module from stage-coupled gene module. We found there is a permissive module with a stage-coupled activation pattern besides the key *Brachyury* module determining notochord fate. The permissive module could be preparing the restriction of notochord fate. Thus, in this case, we show convergent differentiation is not simply driven by a core fate-restricted program (like *Brachyury* module for notochord). Multiple gene modules were also found in the convergent differentiation of myogenesis in mouse embryo (Cao et al., 2019b). However, without the clear stage information we have in *Ciona*, it is not feasible to infer their relationship.

## Acknowledgments

We thank Drs. Kathryn Anderson, Mary Baylies, Alexandra Joyner, Anna-Katerina Hadjantonakis, Danwei Huangfu and Elizabeth Lacy for discussions. ZB is partially supported by an NIH grant (R01GM097576) and an NIH center grant to MSKCC (P30CA008748). WS is supported by the Funding Project of National Key Research and Development Program of China (2018YFD0900604), Natural Science Foundation of China (41676119, 41476120) and start-up fund from Ocean University of China. TY is partially funded by NIH (R01GM124061, R37AI051231 and U19AI057266). BD is supported by the Fundamental Research Funds for the Central Universities from the Ocean University of China (201822016).

## Author contributions

WS, TY and ZB conceived the project. TZ and WS performed the single cell sequencing experiments. TZ, YX, TF, TY and ZB performed the bioinformatics analysis. KI and YS performed the *in situ* experiments. WS and ZB wrote the paper with input from all authors.

## Methods

### Animals and embryology

Fecund *C. savignyi* animals were collected from the Jiaozhou Bay in Qingdao, Shandong and kept in 18°C circulation sea water tank. Fertilization, dechorionation and embryo cultures were done as previously described (Christiaen et al., 2009b). For FGF-MAPK drug inhibitor experiments, embryos were grown in sea water containing 2ug/ml U0126.

For embryo dissociation, staged embryos were transferred to Ca^2+^/Mg^2+^ free artificial sea water (Christiaen et al., 2009b) containing freshly prepared 0.1% trypsin (MP Biomedicals). Single embryos were gently pippeted using a mouth pippet until all blastomeres dissociated. All blastomeres from a single embryo were transferred to a new dish of fresh sea water at 4°C, after which cells were individually transferred using a glass capillary needle attached to a mouth pippet to PCR tubes containing the Smart-seq2 lysis buffer (Picelli et al., 2014). The tubes were flash frozen in liquid nitrogen and temporarily stored at −80°C before library preparation.

### Single-cell RNA library preparation and sequencing

A modified Smart-seq2 protocol with 8bp barcode and 9bp UMI in RT-primers was used to allow sample pooling and amplification bias removing. Reverse transcription and pre-amplification were processed as the Smart-seq2 protocol. Briefly, cells in lysis buffer were thawed on ice, put on 72 °C thermal cycler for 3 min to denature and immediately put on ice for annealing. RT mixture with LNA TSO primer was added and proceeded using the following PCR steps: 42 °C for 90 min, then ten cycles of: 50 °C for 2min, 42 °C for 2min, and finally, 70 °C 15min. cDNA pre-amplification was performed for 12, 13, 14, 15, 16 and 17 cycles respectively for 1/2-cell, 4-cell, 8-cell, 16-cell, 32-cell and 64/110-cell stage samples.

cDNA libraries from different cells were pooled and purified for subsequent sequencing library construction. 500 pg cDNA of each pooled sample was fragmented, tagged, and the 3’end amplified according to the Nextera XT instructions except that a custom P5 primer was used for amplification. Libraries were sequenced on Illumina HiSeq X10 platform with a custom read1 sequencing primer. We used pair-end 150bp and an 8bp index read, consistent with default X10 machine settings. Each cell was sequenced for an average depth of 500,000 paired-end reads.

### In situ hybridization

We performed *in situ* hybridization in both wild-type and U0126-treated embryos at the 64-cell stage. Embryos were hybridized *in situ* with probes for *ZicL* (Imai et al., 2002b) and *lefty* (Imai, 2003) using standard protocols described previously (Yasuo and Satoh, 1994).

### Gene classification

1. Ortholog mapping between *C. savignyi* and *C. intestinalis* *Ciona savignyi* gene model (CSAV2.0; Ensembl) was first compared against the gene models from *Ciona intestinalis* (KH2012; Aniseed) by BLASTP in order to transfer the functional annotation of the genes in *C. intestinalis* by orthology. Mutual best hit with identity > 30% and e-value < 1e-3 were considered as orthologs.
2. Transcription factors (TFs) TFs (n = 387) in *Ciona savignyi* were downloaded from DBD (database of predicted transcription factors) (Charoensawan et al., 2010). TFs (n = 1506) in *Mus musculus* were downloaded from TcoF-DB v2 (Schmeier et al., 2017).
3. Ortholog mapping between *C. savignyi* and mouse *Ciona* and mouse orthologous genes were downloaded from Ensembl. In addition, DEGs in *Ciona* that do not have Ensembl orthologs were matched to mouse genes based on sequence similarity. Specifically, top BLASTP hits (maximal n = 3) with e-value < 10^−10^ and alignment length > 30% were considered as orthologs. Among the DEGs in *Ciona*, 158 genes have Ensembl-assigned orthologs and 45 have orthologs upon further mapping (Table S7).
4. Gene Ontology (GO) term GO annotations for each gene were downloaded from Ensembl. Thirteen categories of GO terms were sequentially matched to GO terms of each DEG in the following order, transcription factor, chromatin, RNA binding/splicing, protease/ubiquitin, signaling pathway, adhesion, extracellular matrix, cytoskeleton/microtubule, myosin/kinesin/dynein, vesicle, mitochondrion, metabolism, and membrane. Each gene is only assigned to one category. Genes whose GO annotation do not matched to any category were assigned to “other”. Genes without any GO annotation were assigned to “unknown”.

### Data processing of scRNA-seq

1. Reads alignment Drop-seq software (Macosko et al., 2015) was used for data de-multiplexing, reads alignment and cellular-molecular barcodes processing. Barcodes (9-16bp) and UMI (17-25bp) extracted from read1 were added to read2 as tags. PolyA tail sequence and template switch oligo sequence were trimmed. Only read2 were mapped to the *C. savignyi* genome (CSAV2.0; Ensembl). A “GE” tag was added onto reads when the read overlaps the exon of a gene. Reads of used exact barcodes were counted to get the raw count matrix. The raw and processed data were deposited into GEO (GSE113788).
2. Filtering of low-quality results Low-quality cells with number of detected genes less than 2000 were removed. Genes with per-embryo effect were considered systemic errors and removed from downstream analysis. Per-embryo effect was defined as greater than 2-fold difference on average UMIs between any pair of embryos at the same developmental stage.
3. Normalization of gene expression Normalization were performed by DESeq2 (Anders and Huber, 2010) in cells at each developmental stage.

### Definition of clusters and DEGs through iterative clustering

1. Selection of HVGs (highly variable genes) We used two methods to select HVGs at each developmental stage as described below. The hypothesis is that HVGs do not conform Poisson distribution as the majority of genes. Genes from both methods were then merged as the HVGs at each stage.
  1.1 Method 1 (Figure S2a).
    1.1.1 Theory: 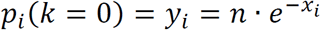 in Poisson distribution, x: mean, y: number of 0-count
    1.1.2 Plot 0-count against log2 mean for each gene
    1.1.3 Tentative outliers on this plot were detected by k-nearest-neighbors distance (R package DDoutlier, k=3-6, distance < 15). The boundary between inliers and outliers was further smoothed by local polynomial regression fitting on maximal inliers on each value on y-axis. Final outliers were considered as genes out of the smooth boundary.
    1.1.4 To exclude genes with low expression, HVGs were defined as outliers with mean > 0.1 and number of 0-count > 10%×n, where n is total number of cells at each stage.
  1.2 Method 2 (Figure S2b).
    1.2.1 Theory: 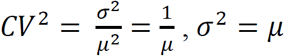 in Poisson distribution
    1.2.2 Plot CV^2^ against log2 mean
    1.2.3 Outliers on this plot were detected by confidence line (p=0.999) of linear regression.
    1.2.4 Same with 1.1.4, to exclude genes with low expression, HVGs were defined as outliers with mean > 0.1 and number of 0-count > 10%×n, where n is total number of cells at each stage.
2. Initial clustering of cells
  2.1 Dimension reduction Dimension reduction at each developmental stage was performed by t-SNE using HVGs (R package Seurat, parameters were set according to cell number, as listed below).

  2.2 Clustering Clustering was performed in t-SNE space with local density clustering (R package dbscan). To optimize parameters of dbscan, we enumerate the combination of parameter *k* (minimal number of points required to form a dense region: 2-6) and parameter *eps* (size of neighborhood: range from the minimal pairwise distance to the maximum pairwise distance in a step of 0.1). The objective function for the parameter search is to minimize the global Davies–Bouldin index 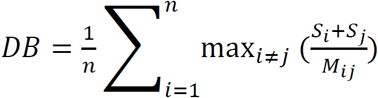, where *S*_*i*_ is the average distance of all points in cluster i to its centroid, *M*_*ij*_ is the distance between the centroids of cluster *i* and cluster *j*, and *n* is the number of clusters. The table below lists the optimized parameters of dbscan and DB index of clustering.

| number of cells | (0,10] | (10,20] | (20,40] | (40,60] | (60,100] | n>100 |
| --- | --- | --- | --- | --- | --- | --- |
| PCA dim | 5 | 10 | 10 | 15 | 20 | 30 |
| t-SNE dim | 5 | 5 | 8 | 10 | 15 | 20 |
| perplexity | 1 | 3 | 5 | 10 | 10 | 20 |

3. Iterative clustering of cells To obtain high resolution on cell types, we applied iterative clustering to resolve nested cell types (Figure S3).
  3.1 Further division of a given cluster
    3.1.1 Clustering of cells is based on the same t-SNE and dbscan-based method. HVGs used are updated to avoid technical noise by filtering HVGs with average UMI > 0.5 in the starting cluster.
    3.1.2 Each tentative cluster produced is evaluated for the difference to its closest cluster, and will only be accepted as a cluster if it is significantly different. The closest cluster is defined by the local Davies–Bouldin index, i.e., cluster *j* in argmax 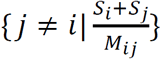 is the closest cluster for cluster *i*. To evaluate the difference between cluster *i* and *j*, we first identify genes from the HVGs whose expression is significantly different between the two groups of cells. For each gene, a p-value is calculated based on the Wilcoxon rank-sum test, where cells are ranked by the UMI of the given gene. A gene is considered significantly different if *p* < 0.01. The cumulative difference for two clusters, d_*ij*_, is defined as 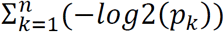, where *p*_*k*_ is the *p* value of significant gene *k* and *n* is the number of significant genes. The significance of d_*ij*_ is further evaluated by a background distribution. This background is calculated by randomly sampling two groups of cells from the whole dataset, with one group matching the number of cells in cluster *i* and the other cluster *j*. A d value is calculated for the two groups. The distribution of d is compiled by repeating the random sampling 1000 times.
    3.1.3 Iterative clustering was applied on each cluster until it reached any of the following 4 terminal conditions. 1) Number of cells less than 4; 2) no partition by dbscan; 3) dbscan identified a sub-cluster from one embryo (residual per-embryo effect as systemic errors in sequencing becomes locally dominant); 4) significance of difference between the tentative clusters show *p* > 0.01 as calculated above.
  3.2 Merge clusters
    3.2.1 Check whether any cluster can be merged after initial clustering, as well as after all clusters meet the termination condition. Merge clusters to its closest cluster if their significance of difference > 0.01.
4. Definition of tentative DEGs (differentially expressed genes) The expression of DEGs has statistical difference between some cell types with high expression and other cell types with low/no expression. Wilcoxon test was used to reveal the statistical difference.
  4.1 Method
    4.1.1 For each gene, sort cell types by mean expression in each type.
    4.1.2 To define high and low/no groups, we divided the sorted list of cell types into two groups at every split between two adjacent types and calculate their significance by Wilcoxon test. The split with minimal p value was used to define high and low/no group
  4.2 Thresholds
    4.2.1 Mean count in high expression group > 0.5
    4.2.2 Fold difference between high and low/no group > 1.8
    4.2.3 Wilcoxon p value cutoffs at all stages (higher cell number gives more significant p value, see below)

| Stage | $k$ | $eps$ | DB index |
| --- | --- | --- | --- |
| C4 | 3 | 38.8 | 0.31 |
| C8 | 4 | 39.3 | 0.38 |
| C16 | 4 | 13.8 | 0.52 |
| C32 | 3 | 4.2 | 0.38 |
| C64 | 4 | 1.8 | 0.50 |
| C110 | 2 | 1.6 | 0.59 |

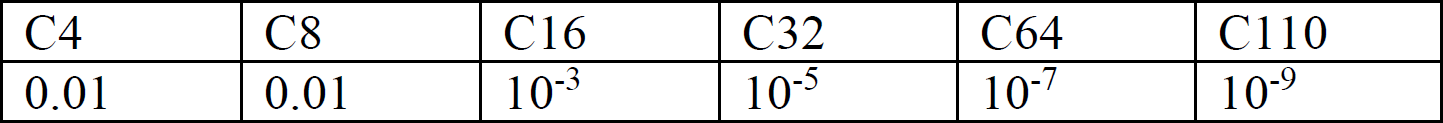
  4.3 Special case
    4.3.1 C16 has the lowest detection ratio of known markers, which may indicate its sequencing quality is relatively low. So we did manual curation on tentative DEGs at C16 defined as above. The idea is to nucleate groups of expression patterns with confident DEGs. First, tentative DEGs with Wilcoxon p value less than 10^−5^ were kept as confident DEGs. Second, for other tentative DEGs, genes whose expression pattern in cell types do not match expression pattern of any confident DEG were filtered out.
5. Reclassification of individual cells (Figure S3) To account for the classification of each sequenced cell, we reclassified the cell type of each cell by tentative DEGs after iterative clustering.
  5.1 Method
    5.1.1 Calculate mean expression of tentative DEGs in each cell type.
    5.1.2 To measure the similarity between an individual cell and a given cell type, Pearson’s correlation was calculated based on genes with non-0 expression (UMI in individual cell > 4 or UMI in cell type > 0.5) and p value of correlation was calculated using Student’s t-distribution with degrees of freedom n-2 (n is the length of vector).
    5.1.3 Assign the cell type with p value less than 10^−5^ as the identity of individual cell. If there are more than one cell type with p value < 10^−5^, assign the cell type with minimal p value.
  5.2 Reject vague cells
    5.2.1 For each cell type in each embryo, sort cells by p value per cell type per embryo and remove cells with large p value. The p value cutoff was defined as the p value of cell that has a 50-fold or greater increase compared to the p value of next cell on the sorted list.
6. Final list of cell types and DEGs After re-assignment, we updated the list of cells belonging to each cell type (Table S5). Final DEGs (Table S5) were computed based on final cell types using the same method of defining tentative DEGs. DEGs of a given cell type were defined as final DEGs that have this cell type in high expression group. Final cell types were visualized by t-SNE using final DEGs (Figure S4).
7. Assignment of lineage identity Assignment of canonical cell types/lineage identity was based on known markers (Imai et al., 2006). The relationship between cell clusters and canonical cell types was documented in Table S6
  7.1 Special cases
    7.1.1 B at 4-cell stage and B4.1 at 8-cell stage: when we proceeded to assign ID and define DEGs, we found B and B4.1 subdivided into two clusters violating expected cell number per embryo. So we rejected the sub-division and took the super-clusters that correspond to B and B4.1.
    7.1.2 The A- and B-notochord at 110-cell stage showed the weakest separation (p=0.036 between the two groups). The legitimacy of their difference is systematically documented in Figure S13c and related text. ZicL-3 expression was used to assign lineage identity between the two groups (Imai et al., 2006).

### Analysis of scRNA-seq data in drug-treated embryos

Data preprocessing and iterative clustering of scRNA-seq data in drug-treated embryos were performed as described above (Figure 1d).

1. Matching cell types between drug-treated and WT embryo For cell types in drug-treated embryos without fate transform, we considered WT cell type with the most similar expression of DEGs as its cell type. For cell types transforming cell fate, the expression of retained DEGs could be used to infer its presumptive cell type. The numbers of cells per cell type per drug-treated embryo agree with expected number of cells.
  1.1 Method
    1.1.1 Calculate non-zero (mean of count > 0.5) correlation of mean expression of DEGs at 64-cell stage between clusters of drug-treated embryos and WT cell types (Figure S14b).
    1.1.2 The most similar cell type in WT type is considered as the cell type for each cell cluster of drug treated embryos.
    1.1.3 DEGs at 64-cell stage were used to distinguish cell types in a mixture cluster caused by fate transformation (Figure S14b).
  1.2 B7.5 fate transformation
    1.2.1 B7.4/5 is a mixture of B7.4 and B7.5. No retained DEG could distinguish B7.5 from B7.4. Because the probability of losing all 4 B7.5 cells is very low (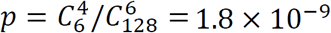, 6 cells were lost in total 128 cells from two drug-treated embryos), we concluded that B7.5 is transformed to B7.4.
2. The global hierarchy of cell type similarities The similarity between cell types in WT and drug treated embryos was visualized by SplitsTree (Huson and Bryant, 2006). As input for SplitsTree, pairwise distances between cell types used in SplitsTree were computed as 1-Jaccard index of the overlap of their DEGs.

### MT co-segregating genes

To search for genes that have similar asymmetric expression as mitochondrion-encoded genes, we required uneven gene expression in all three pairs of asymmetrical divisions (A7.4/8 > A7.3/7, B7.4 > B7.3, and B7.8 > B7.7). Specifically, in WT embryos, genes with fold differences in three pairs greater than 1.5 and average UMI greater than 0.5 in high-expression cells (A7.4/8, B7.4, and B7.8) were defined as mitochondrion co-segregating genes. In drug treated embryos, mitochondrion co-segregating genes with 1.5 or higher fold differences in A7.4/8 vs. A7.3/7 and B7.4/8 vs. B7.3/7 were considered as remaining asymmetrical, and with 1.2 or lower fold differences in those two pairs were considered as losing asymmetrical.

### Comparison of cell types between Ciona and mouse

We measured the similarity of cell types between species by shared DEGs, especially TFs.

1.1 Cell types and their DEGs
  1.1.1 Cell types at 110-cell stage in *Ciona* were used except B7.6 and epidermis, which do not have homologous cell type in mouse E8.25 embryos. DEGs of close-related cell types were merged, i.e., merging DEGs of A8.5/6/13/14 and B8.6 as notochord, A8.7/8 and A8.15/16 as nerve cord.
  1.1.2 Cell types in E8.25 mouse embryos (Ibarra-Soria et al., 2018) were used except blood, endothelial, and placodes. DEGs of mouse cell types were obtained from the accompanying database (https://marionilab.cruk.cam.ac.uk/organogenesis/) and further filtered with our thresholds in *Ciona*, i.e., fold difference > 1.8 and mean of count > 0.5.
1.2 Score similarity
  1.2.1 The sum of scores from overlapped genes is the score of pair of cell types between mouse and *Ciona*. The score of individual overlapped gene is given as following, Specific TF (# of expressed *Ciona* cell types < 2): 6; TF (2 <= # of expressed *Ciona* cell types < 5): 4; TF (5 <= # of expressed *Ciona* cell types): 3; Muscle effector genes: 1; Other genes: 0.5.
  1.2.2 Match cell types between mouse and *Ciona* based on scores. The principle of threshold design is that cell type pair with high similarity have a score equivalent to score of two or more TFs. High similarity is defined as, for each mouse cell type, the best matched *Ciona* cell type has a score > 10. Moderate similarity is defined as, for each mouse cell type, the matched *Ciona* cell types have a score > 10 and the difference to best score <= 4.

## Supplementary Materials

This contains all the legends for Supplementary Tables (S1-S7) and Figures (S1-S15). Table S1-S3 are provided in pdf format.

## Supplementary Tables

Table S1. Number of expected, lost, low quality and unclassified cells per embryo per stage.

Table S2. Number of detected (numerators) vs. expected (denominators) cells for each identified cell type across embryonic stages.

Table S3. Examination of unresolved cell types.

Table S4. Cell_type_average_expression.xlsx

Table S5. Cell_type_DEGs.xlsx

Table S6. Cell_cluster_to_lineage.xlsx

Table S7. Ciona_Mouse_comparison.xlsx

## Supplementary Figures

Figure S1. The cell lineage of the early Ciona embryo with blastomere names.

Figure S2. Detection of highly variable genes (HVGs).

Figure S3. Computational analysis at the 64-cell stage to define cell types and classify individual cells.

Figure S4. Cell clustering at each stage.

Figure S5. Comparison of Levine’s cell types at 110-cell stage.

Figure S6. Quality control of identified DEGs.

Figure S7. List of DEGs at the 4-, 8-, 16- and 32-cell stage.

Figure S8. List of DEGs at the 64-cell stage.

Figure S9. List of DEGs at the 110-cell stage.

Figure S10. Comparison of expression sites for known cell type-specific marker genes between published in situ results in C. intestinalis (Imai et al., 2006) and detection in this study.

Figure S11. Prediction of zygotic expression.

Figure S12. Enrichment analysis of known *postplasmic/PEM* genes.

Figure S13. Gene expression dynamics in notochord lineage differentiation.

Figure S14. Fate transformations and asymmetry of MT genes upon U0126 treatment.

Figure S15. Comparison of DEG expression between wild-type and U0126-treated embryos.

**Table S1.** Number of expected, lost, low quality and unclassified cells per embryo per stage.

| Stage | Total high quality | Embryo | Lost | Low-quality | Unclassified cells |
| --- | --- | --- | --- | --- | --- |
| 1-cell | 8 | E01 | 0 | 0 | - |
|  |  | E02 | 0 | 0 | - |
|  |  | E03 | 0 | 0 | - |
|  |  | E04 | 0 | 0 | - |
|  |  | E05 | 0 | 0 | - |
|  |  | E06 | 0 | 0 | - |
|  |  | E07 | 0 | 0 | - |
|  |  | E08 | 0 | 0 | - |
| 2-cell | 10 | E01 | 0 | 0 | - |
|  |  | E02 | 0 | 0 | - |
|  |  | E03 | 0 | 0 | - |
|  |  | E04 | 0 | 0 | - |
|  |  | E05 | 0 | 0 | - |
| 4-cell | 12 | E01 | 0 | 0 | 0 |
|  |  | E02 | 0 | 0 | 0 |
|  |  | E03 | 0 | 0 | 0 |
| 8-cell | 24 | E01 | 0 | 0 | 0 |
|  |  | E02 | 0 | 0 | 0 |
|  |  | E03 | 0 | 0 | 0 |
| 16-cell | 32 | E01 | 0 | 0 | 2 |
|  |  | E02 | 0 | 0 | 0 |
| 32-cell | 65 | E01 | -2* | 1 | 1 |
|  |  | E02 | 0 | 0 | 0 |
| 64-cell | 173 | E01 | 2 | 2 | 3 |
|  |  | E02 | 1 | 2 | 2 |
|  |  | E03 | 10 | 2 | 1 |
| 110-cell | 304 | E01 | 10 | 3 | 5 |
|  |  | E02 | 0 | 9 | 7 |
|  |  | E03 | 3 | 1 | 2 |
| Drug 64-cell | 122 | E01 | 0 | 1 | 1 |
|  |  | E02 | 3 | 2 | 1 |
| Total | 750 | 31 | 29 | 23 | 25 |

\*Harvested 2 more cells than expected at 32-cell stage

**Table S2.** Number of detected (numerators) vs. expected (denominators) cells for each identified cell type across embryonic stages.

| Stage | No. of embryos | No. of cell types<br>(detected/known)<br>-cell types mixed together | No. of cells per type<br>(cell types--detected/expected) |  |  |  |
| --- | --- | --- | --- | --- | --- | --- |
| 1-cell | 8 | **1/1** | **Total: 8/8** |  |  |  |
| 2-cell | 5 | **1/1** | **Total: 10/10** |  |  |  |
| 4-cell | 3 | **2/2** | **Total: 12/12** |  |  |  |
|  |  |  | A | 6/6 |  |  |
|  |  |  | B | 6/6 |  |  |
| 8-cell | 3 | **2/4** | **Total: 24/24** |  |  |  |
|  |  | 1) A4.1,a4.2,B4.2 (A4.1; a4.2; B4.2) | A4.1,a4.2,B4.2 | 18/18 |  |  |
|  |  |  | B4.1 | 6/6 |  |  |
| 16-cell | 2 | **4/5** | **Total: 30/32** |  |  |  |
|  |  | 1) animal (a5.3/4; b5.3/4) | A5.1/2 | 6/8 |  |  |
|  |  |  | animal | 16/16 |  |  |
|  |  |  | B5.1 | 4/4 |  |  |
|  |  |  | B5.2 | 4/4 |  |  |
| 32-cell | 2 | **7/7** | **Total: 64/64** |  |  |  |
|  |  |  | A6.1/3 | 8/8 |  |  |
|  |  |  | A6.2/4 | 8/8 |  |  |
|  |  |  | Animal | 32/32 |  |  |
|  |  |  | B6.1 | 4/4 |  |  |
|  |  |  | B6.2 | 4/4 |  |  |
|  |  |  | B6.3 | 4/4 |  |  |
|  |  |  | B6.4 | 4/4 |  |  |
| 64-cell | 3 | **14/19** | **Total: 167/192** |  |  |  |
|  |  | 1) A7.4/8 (A7.4; A7.8) | A7.1/2/5 | 18/18 | B7.5 | 3/6 |
|  |  | 2) a7.9/10/13 (a7.9/10; a7.13) | A7.3/7 | 8/12 | B7.6 | 3/6 |
|  |  | 3) epidermis (a7.11/12/15/16; a7.14; b7.11/13/14; b7.12/15/16) | A7.4/8 | 11/12 | B7.7 | 5/6 |
|  |  |  | A7.6 | 6/6 | B7.8 | 6/6 |
|  |  |  | B7.1/2 | 10/12 | a7.9/10/13 | 13/18 |
|  |  |  | B7.3 | 6/6 | b7.9/10 | 10/12 |
|  |  |  | B7.4 | 6/6 | epidermis | 62/66 |
| 110-cell | 3 | **16/24** | **Total: 290/330** |  |  |  |
|  |  | 1) A8.15/16 (A8.15; A8.16) | endoderm | 28/30 | B7.5 | 5/6 |
|  |  | 2) B8.15/16 (B8.15; B8.16) | A8.5/6/13/14 | 20/24 | B7.6 | 3/6 |
|  |  | 3) anterior neu (a8.17/19; a8.18/20; a8.25; a8.26) | A8.7/8 | 12/12 | B7.7 | 4/6 |
|  |  | 4) posterior neu (b8.17/19; b8.18/20) | A7.6 | 6/6 | B8.15/16 | 12/12 |
|  |  | 5) anterior ep (a8.21/22; a8.23/24/27/28; a8.29/30/31/32) | A8.15/16 | 10/12 | anterior neu | 26/36 |
|  |  |  | B8.5 | 6/6 | posterior neu | 24/24 |
|  |  |  | B8.6 | 4/6 | anterior ep | 53/60 |
|  |  |  | B8.7/8 | 12/12 | posterior ep | 65/72 |

**Table S3.** Examination of unresolved cell types.

| Stage | Cell type | Known sub-types | Genes that are known to have different expression between sub-types | Global high-quality genes | Number of cells that match expression pattern of high-quality genes in sub-types |
| --- | --- | --- | --- | --- | --- |
| C8 | A4.1, a/b4.2 | A4.1<br>a4.2<br>b4.2 | KH.C11.570<br>KH.L7.6<br>KH.C7.226<br>FoxA-a | - | - |
| C16 | a5.3/4,<br>b5.3/4 | a5.3/4<br>b5.3/4 | KH.C11.529<br>KH.C9.541,<br>FoxA-a | - | - |
| C64 | a7.9/10/13 | a7.9/10<br>a7.13 | AP-2-like2,<br>Otx | Otx | - |
| C64 | A7.4/8 | A7.4<br>A7.8 | Irx-B | Irx-B | - |
| C64 | epidermis | a7.11/12/15/16<br>a7.14<br>b7.11/13/14<br>b7.12/15/16 | E(spl)/hairy-a, Emc,<br>SoxB2, C3H | E(spl)/hairy-a, Emc,<br>C3H | 6 out of 62<br>(see below) |
| C110 | A8.15/16 | A8.15<br>A8.16 | E(spl)/hairy-b,<br>Neurogenin | Neurogenin | - |
| C110 | B8.15/16 | B8.15<br>B8.16 | Otx, PPAR,<br>SoxF | - | - |
| C110 | a-line neural cells | a8.17/19<br>a8.18/20<br>a8.25<br>a8.26 | FoxC, Otx,<br>ZicL-3,<br>C3H, AP-2-like2 | ZicL-3 | - |
| C110 | b-line neural cells | b8.17/19<br>b8.18/20 | E(spl)/hairy-a, FoxH-b,<br>RAR,<br>SoxB1,<br>ZicL-3 | FoxH-b,<br>RAR,<br>SoxB1,<br>ZicL-3 | 3 out of 26<br>(see below) |
| C110 | a-line epidermis | a8.21/22<br>a8.23/24/27/28<br>a8.29/30/31/32 | E(spl)/hairy-a, C3H | - | - |
| total | 10 |  | 26 | 10 |  |

\*C8 markers are from Aniseed database \*C16 markers are from Ilsley 2017 bioRxiv

**Figure S1.**
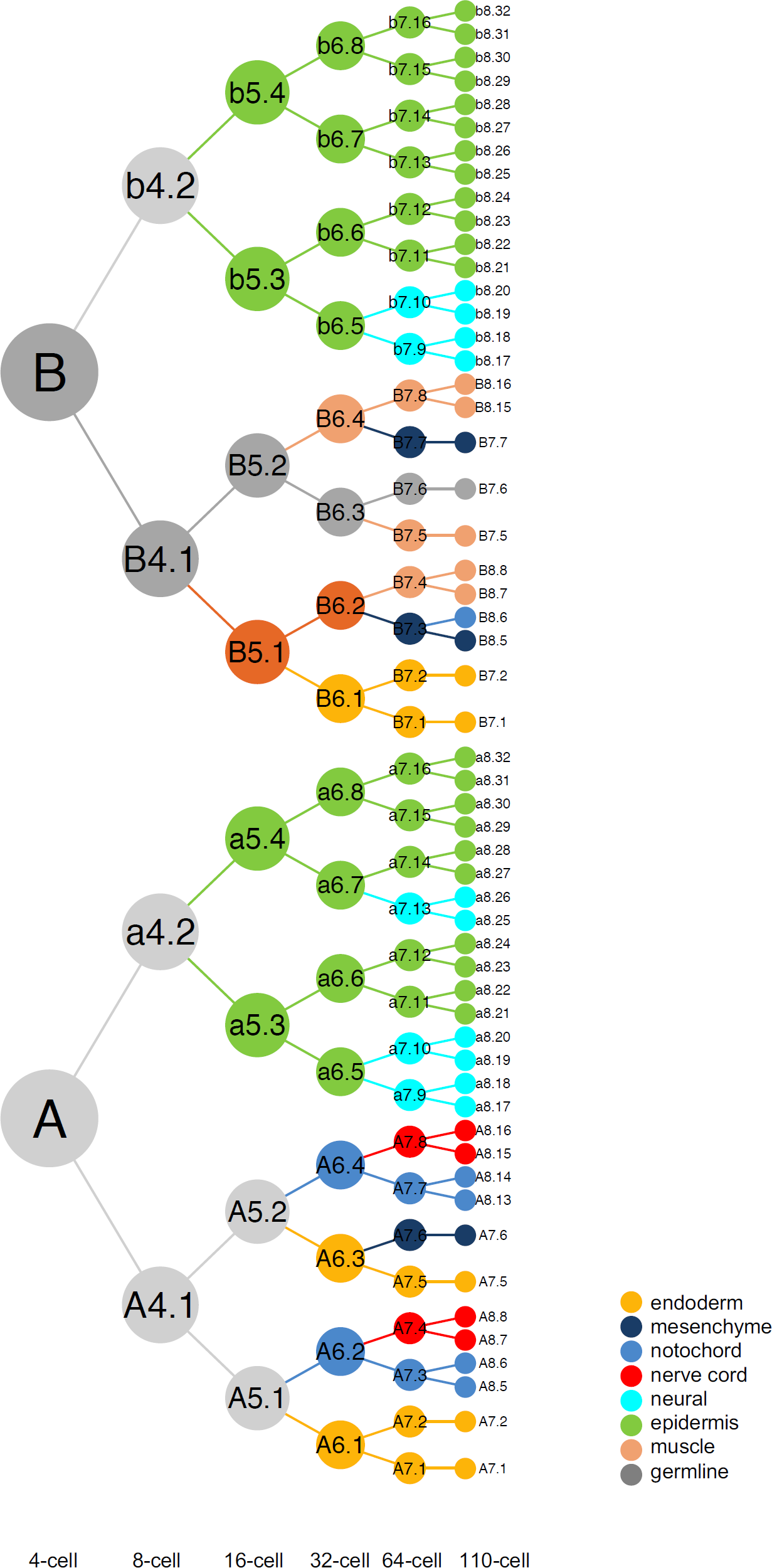
The cell lineage of the early *Ciona* embryo with blastomere names.

**Figure S2.**
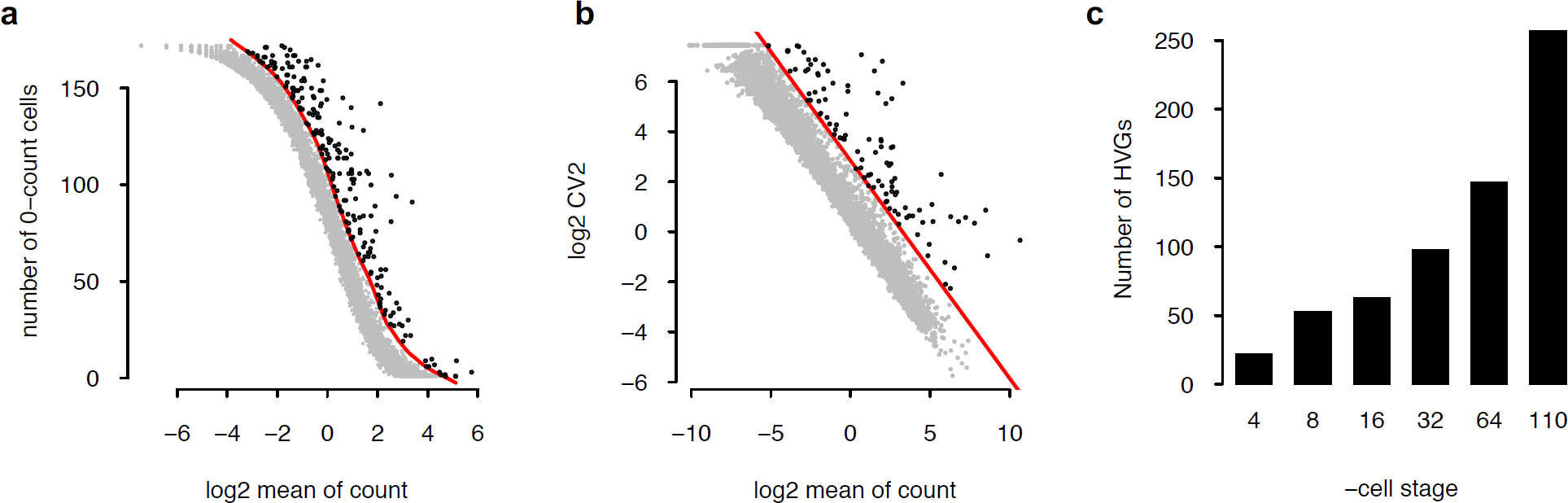
Detection of highly variable genes (HVGs). **(a)** The first filter for HVGs examines the relationship between log_2_ of the mean UMI of a gene across all cells and the number of cells with 0 UMI for each gene. The red curve is calculated based on a chosen level of deviation from a Poisson distribution. Genes (black dots) above the red curve are selected. **(b)** The second filter for HVGs examines the relationship between log_2_ of the mean UMI of a gene across all cells and log_2_ of CV2 of its UMI across all cells for each gene. The red line is calculated based on a chosen level of deviation from a Poisson distribution. Genes (black dots) above the red line are selected. **(c)** Number of HVGs per embryonic stage.

**Figure S3.**
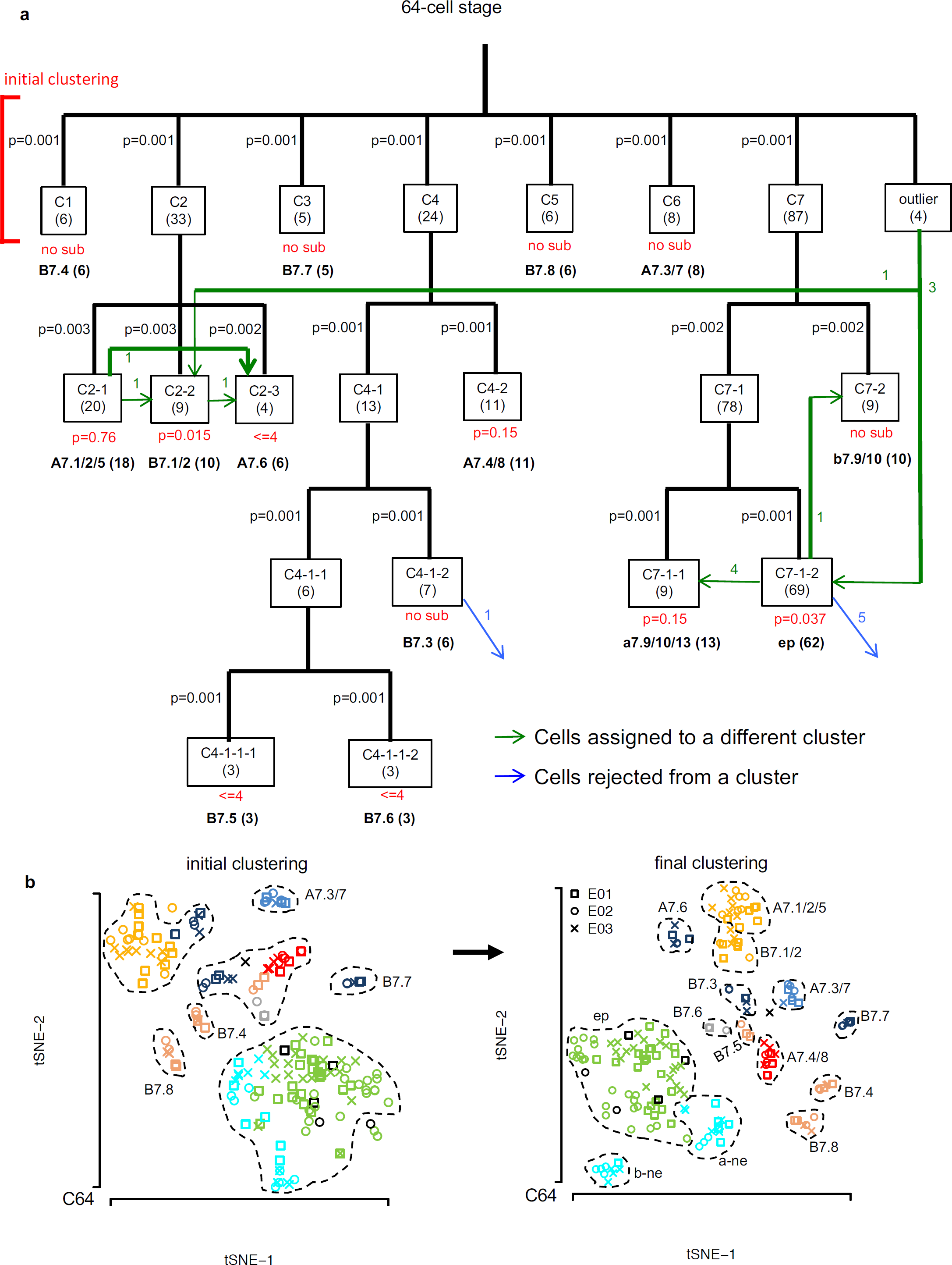
Computational analysis at the 64-cell stage to define cell types and classify individual cells. **(a)** Flow chart showing the results at each step of clustering, iterative clustering and cell classification. Each box represents a cluster of cells (numbered with a C prefix). Outlier is the collection of individual cells that were not assigned to any clusters by DBSCAN. Numbers in parenthesis in each box show the number of cells in the corresponding cluster. p-values in black above each box show the significance of the cluster. Red text below boxes show the reason that the corresponding cluster was not further divided. “<=4” denotes the situation where the number of cells in the cluster <=4 and there too small to be further divided; “no sub” denotes the situation where no division by DBSCAN; p-values denotes the situation where the putative sub-clusters by DBSCAN were not statistically significant (p>=0.01), with the p-value reported. Green lines and associated numbers show the number of cells reassigned from one cluster to another based on DEG similarity. Blue lines and associated numbers show the number cells rejected from a cluster but not re-assigned to any other. Blastomere names assigned at the end are in bold, with the number of cells in the corresponding cell type in parenthesis. ep: epidermis. **(b)** Initial clustering of cells based on the HVGs (left) before iterative clustering and cell classification and the final clusters based on the final defined DEGs (right). Conventions follow Figure 2a.

**Figure S4.**
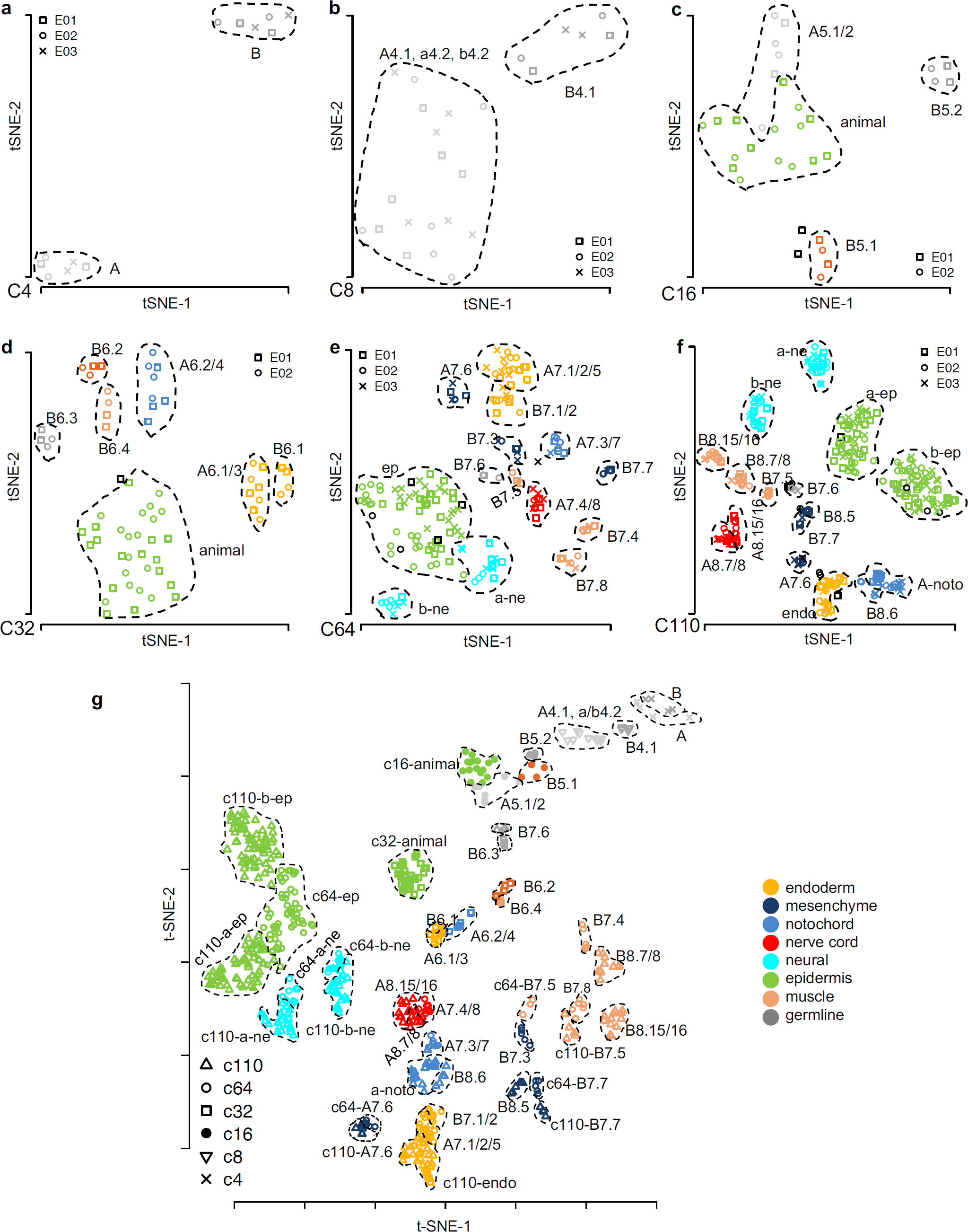
Cell clustering at each stage. **(a-f)** Defined cell clusters at the 4-cell (**a**), 8-cell (**b**), 16-cell (**c**), 32-cell (**d**), 64-cell (**e**) and 110-cell (**f**) stage. Cells from different embryos are represented by different symbols. Black symbols represent rejected/unclassified cells from corresponding embryos. **(g)** Display of cell clusters across all embryonic stages. For visual clarity, cells from the early stages (4-, 8-, 16-cell) are shifted by 28, 16, 4 in t-SNE1 and 18, 16, 12 in t-SNE2, respectively. The coordinates of B-line mesodermal cells at the 64- and 110-cell stages are scaled by 2 in t-SNE1.

**Figure S5.**
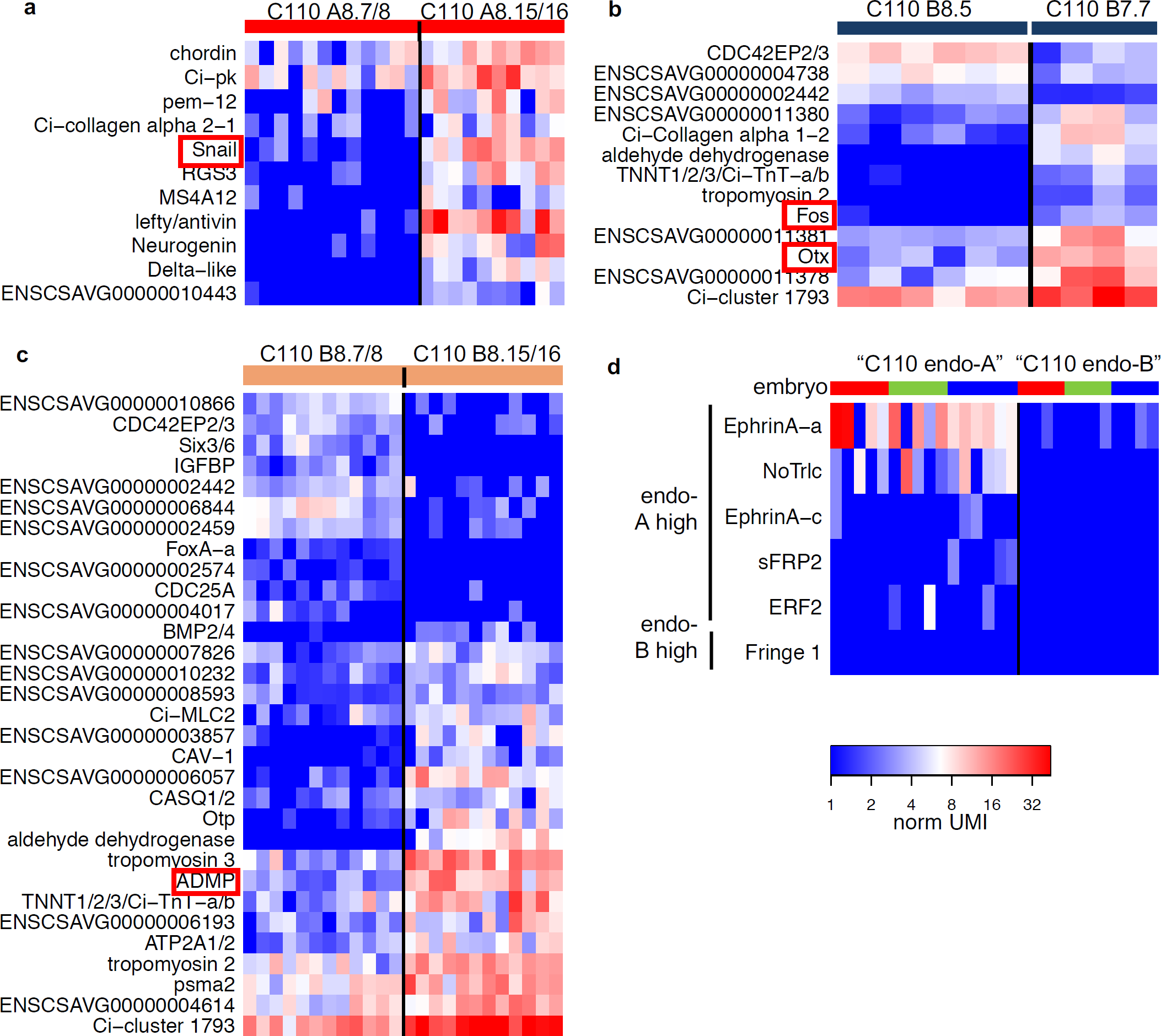
Comparison of Levine’s cell types at 110-cell stage. **(a-c)** For each pair of cell types not resolved in (Cao et al., 2019a), the corresponding heatmap shows DEGs with significantly different expression levels between the two groups of cells (rank statistics p<0.01). Red boxes indicate known *in situ* markers that distinguish the corresponding cell types. **(d)** Manual sorting of endoderm cells based on the expression of six genes reported in (Cao et al., 2019a) to distinguish A-line and B-line endoderm. Red, green and blue bars at the top denote cells from each of the three embryos. These numbers are consistent with the expected numbers, indicating that this sorting may be reasonable.

**Figure S6.**
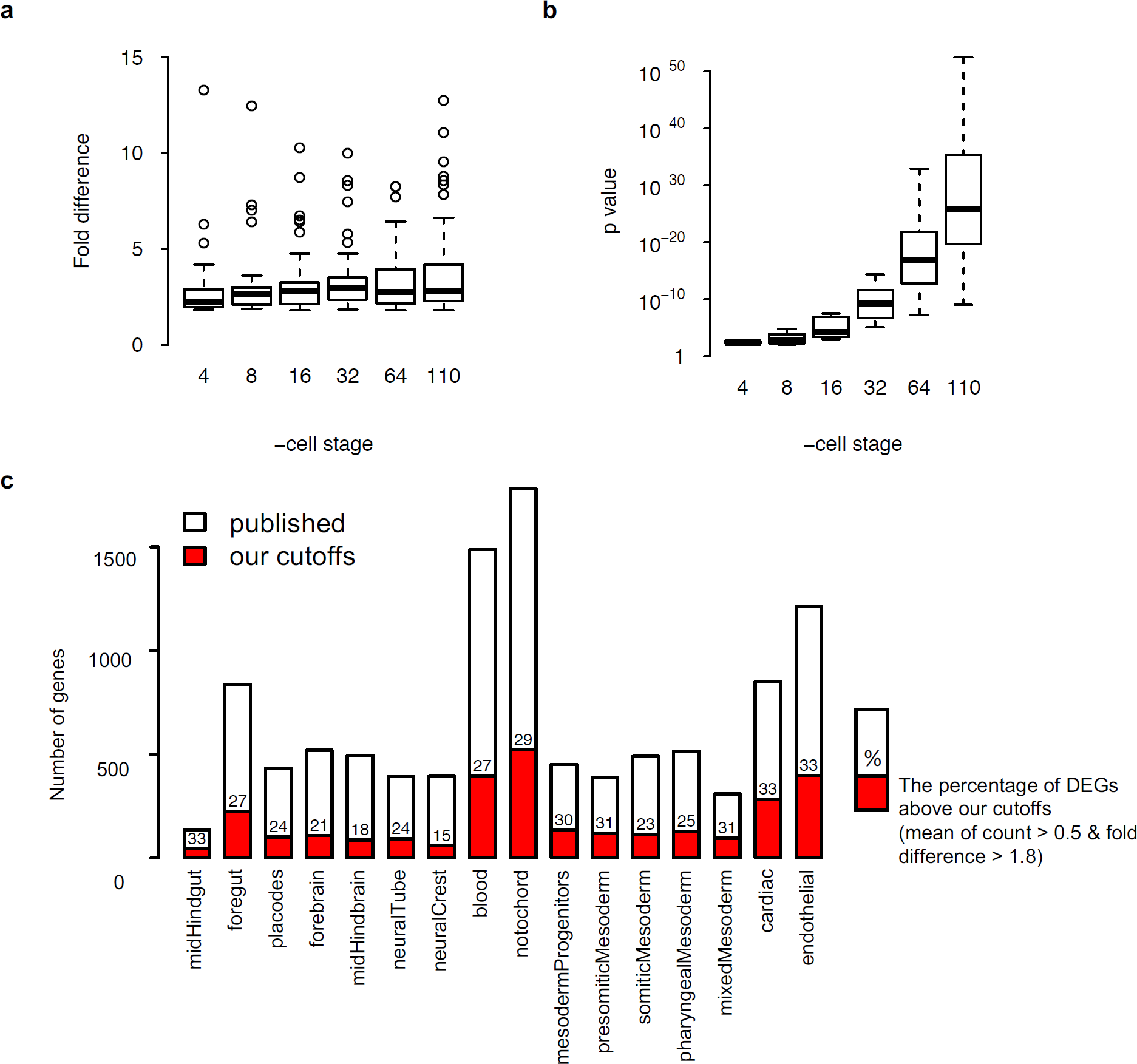
Quality control of identified DEGs. **(a)** Fold difference of DEG expression levels between expressing and low/no expressing cells. Each dot represents a DEG. **(b)** p-value of Wilcoxon rank-sum test of DEGs. **(c)** Filtering of published DEGs for Mouse E8.25 (Ibarra-Soria et al., 2018) with thresholds in this study. Red: DEGs passed the thresholds.

**Figure S7.**
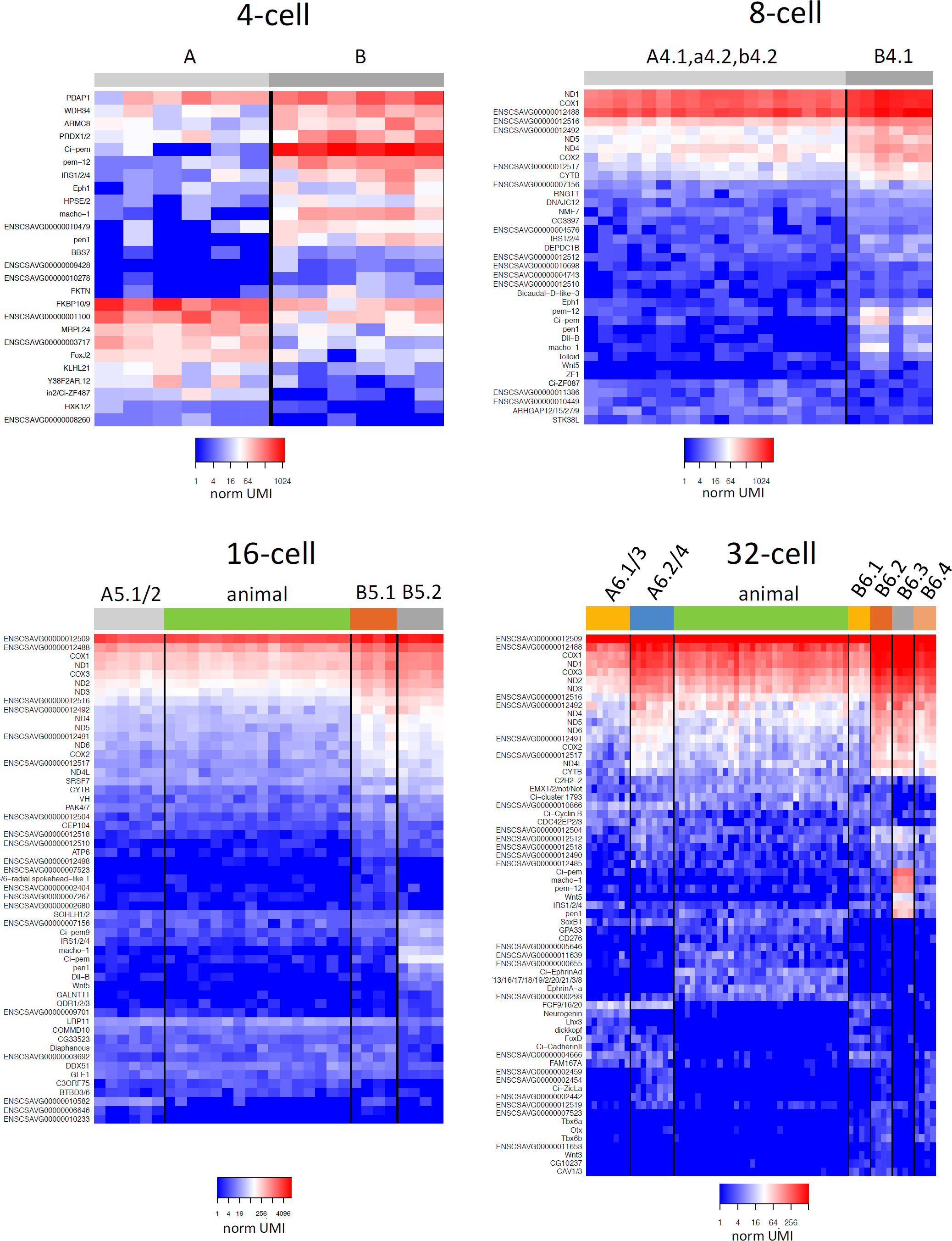
List of DEGs at the 4-, 8-, 16- and 32-cell stage. Heatmap convention follows Figure 2d. See Table S5 for details.

**Figure S8.**
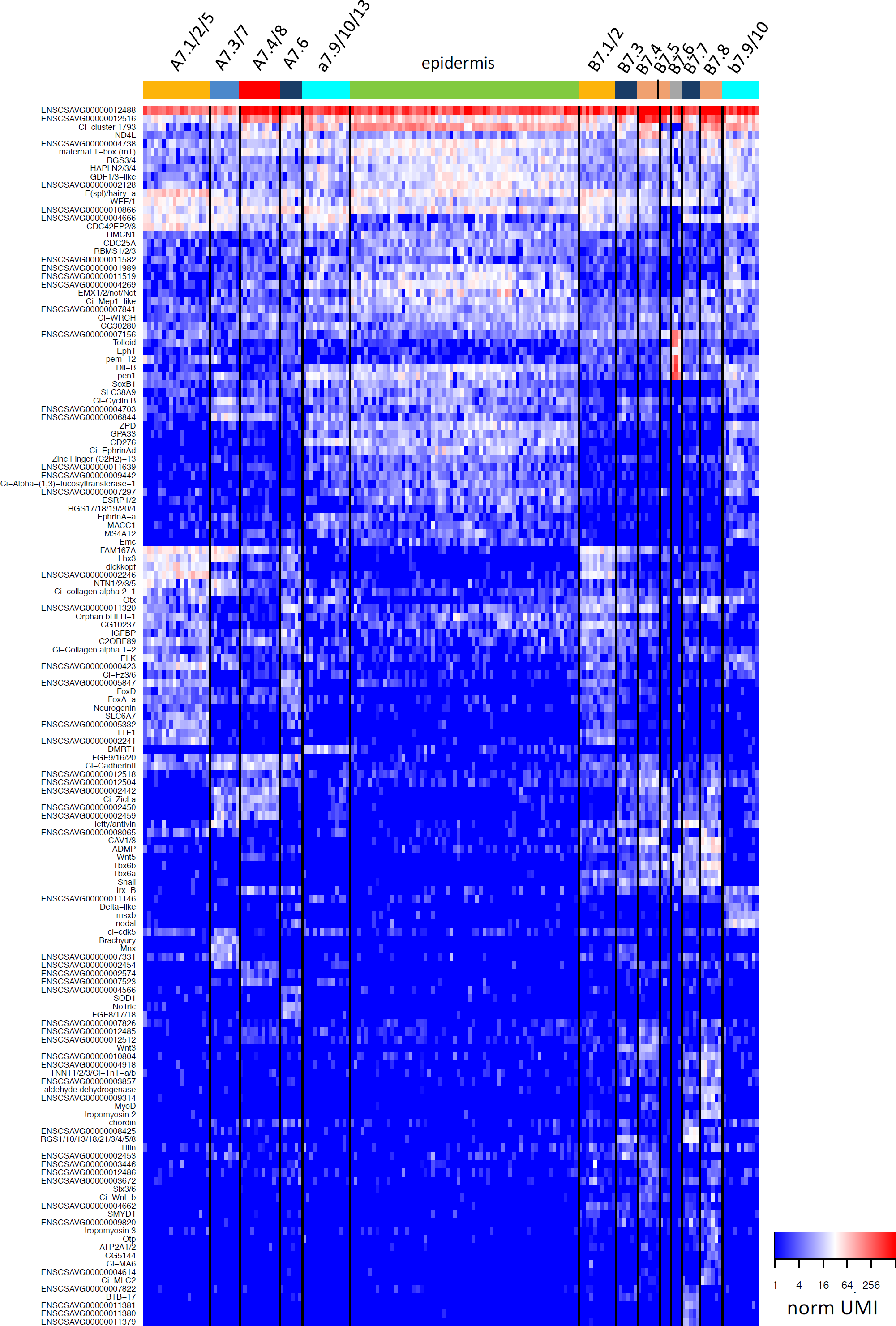
List of DEGs at the 64-cell stage.

**Figure S9.**
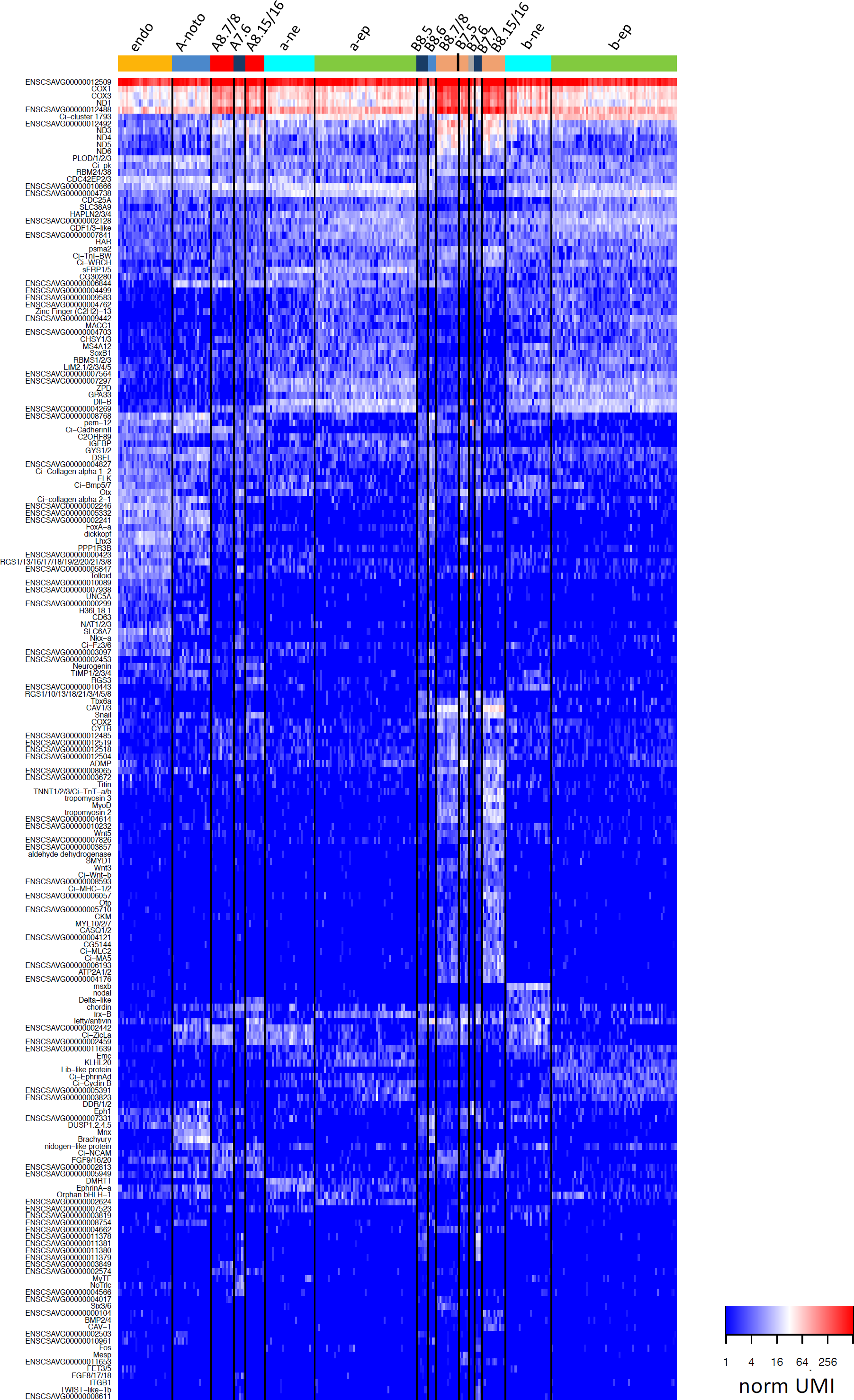
List of DEGs at the 110-cell stage.

**Figure S10.**
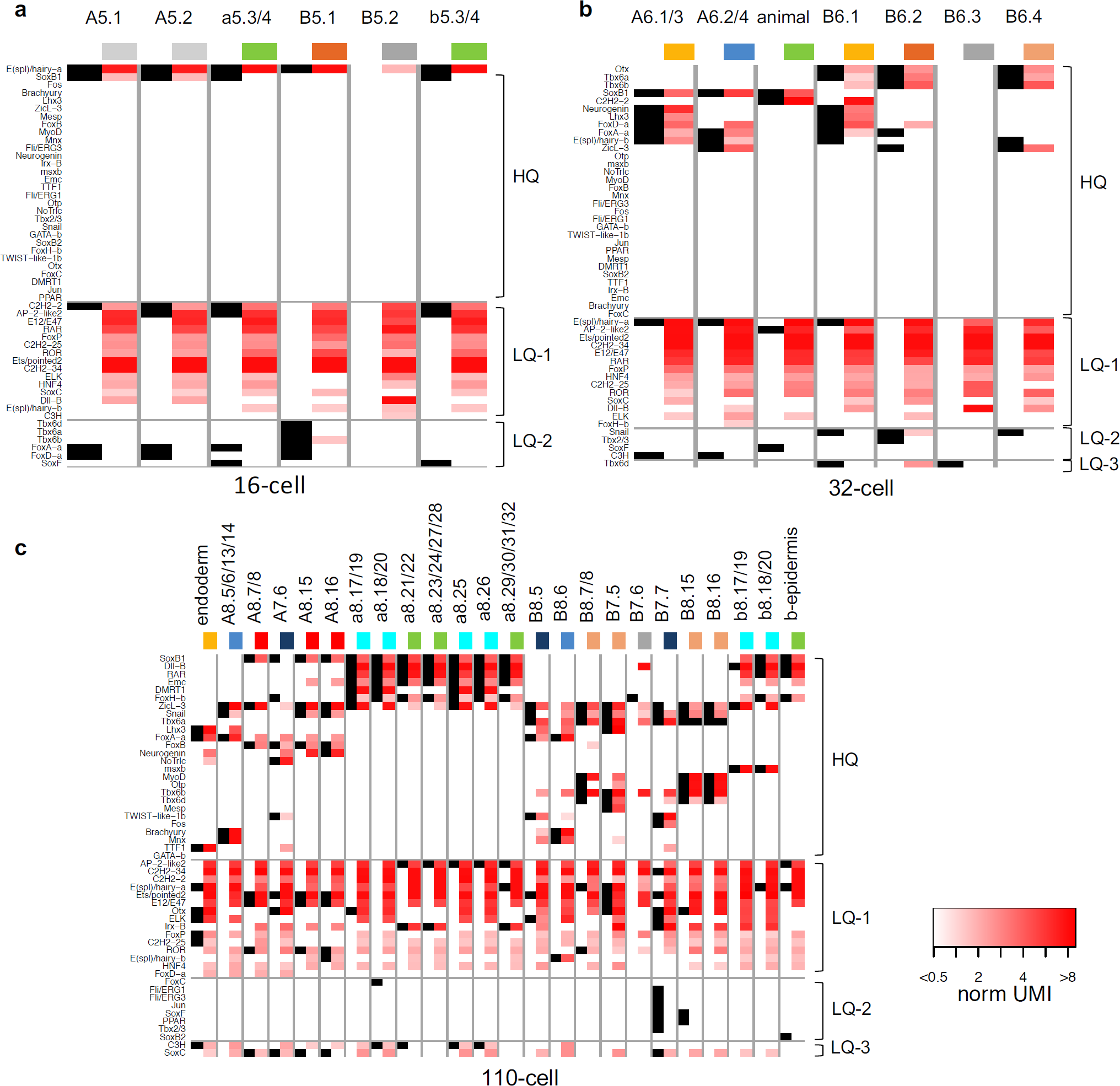
Comparison of expression sites for known cell type-specific marker genes between published in situ results in C. intestinalis (Imai et al., 2006) and detection in this study. **(a-c)** Comparison at the 16-cell (**a**), 32-cell (**b**) and 110-cell (**c**) stage. Convention follows Figure 3g.

**Figure S11.**
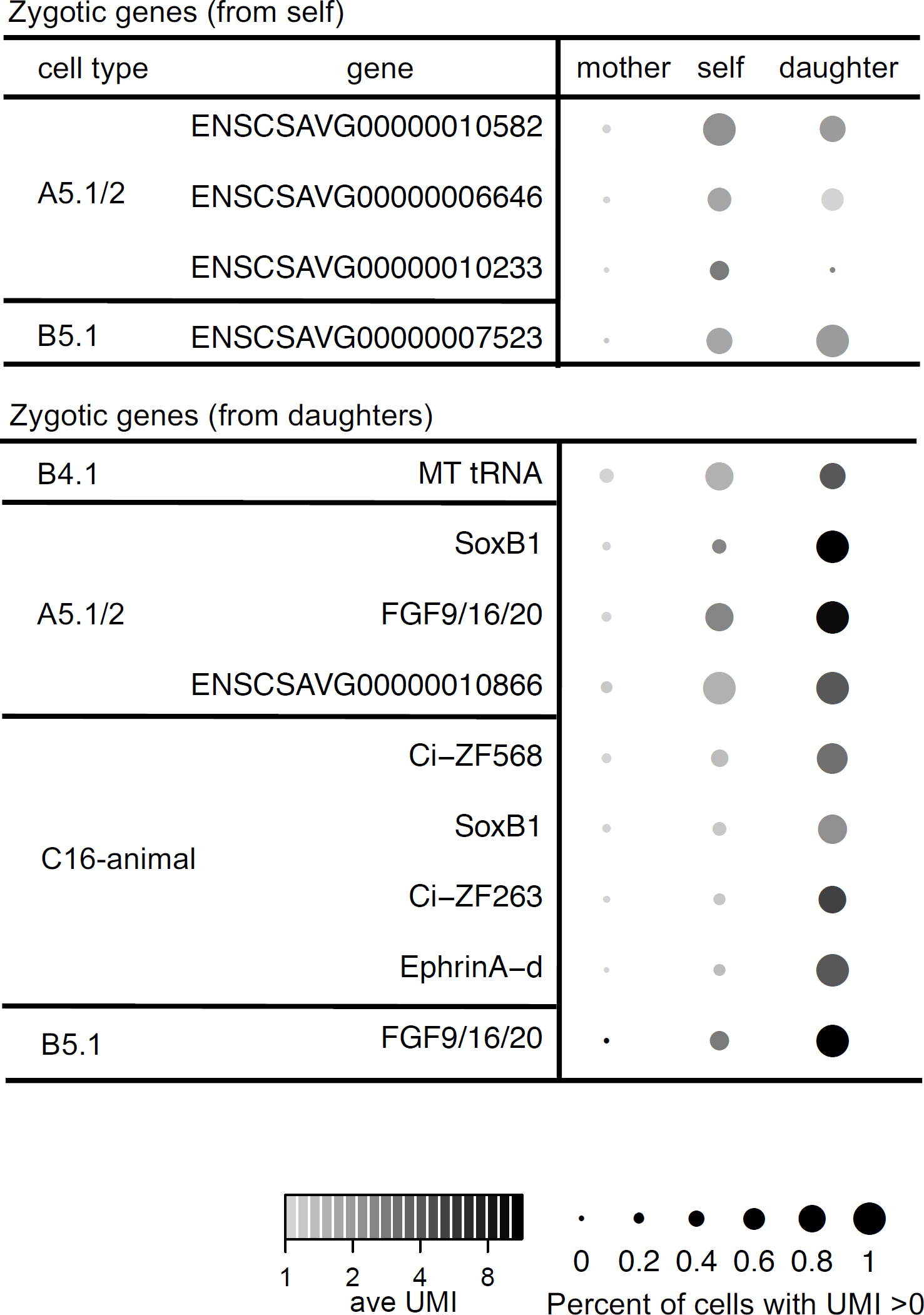
Prediction of zygotic expression. Mother, self and daughter denote the cell named in the first column, its mother cell, and one of its daughter cells with higher average UMI of the corresponding gene. “From self” denote cell types where significant expression was detected in itself. “From daughter” denote cell types where expression was detected in itself and significant expression was detected in its daughter. Circle size shows the fraction of cells in the corresponding type with UMI>0. Shade of color shows average UMI in the type.

**Figure S12.**
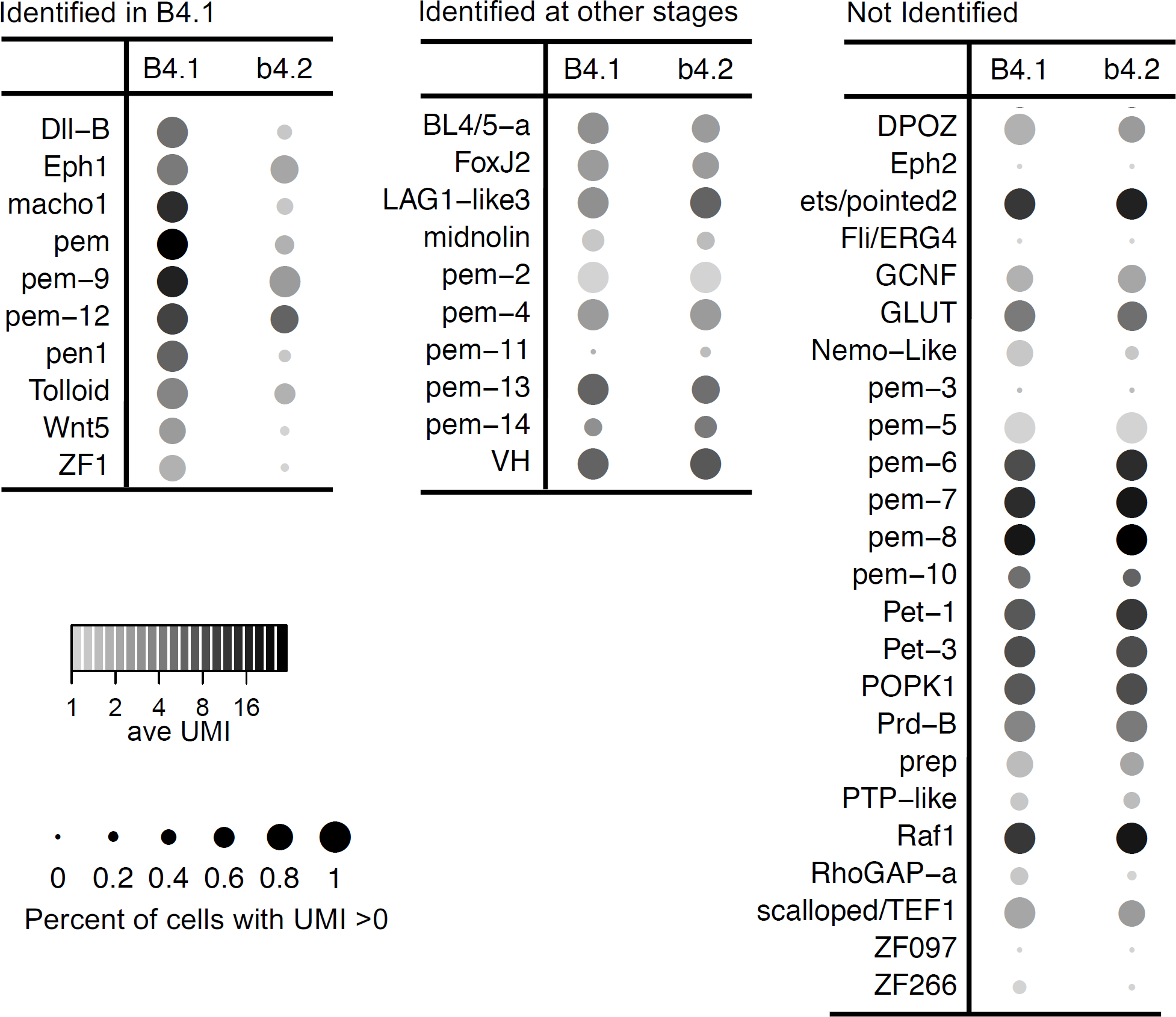
Enrichment analysis of known *postplasmic/PEM* genes. Comparison of PEM genes in B4.1 vs. b4.2 for PEMs enriched in B4.1, enriched in other germline stages or not identified by enrichment analysis. Table convention follows Figure S11.

**Figure S13.**
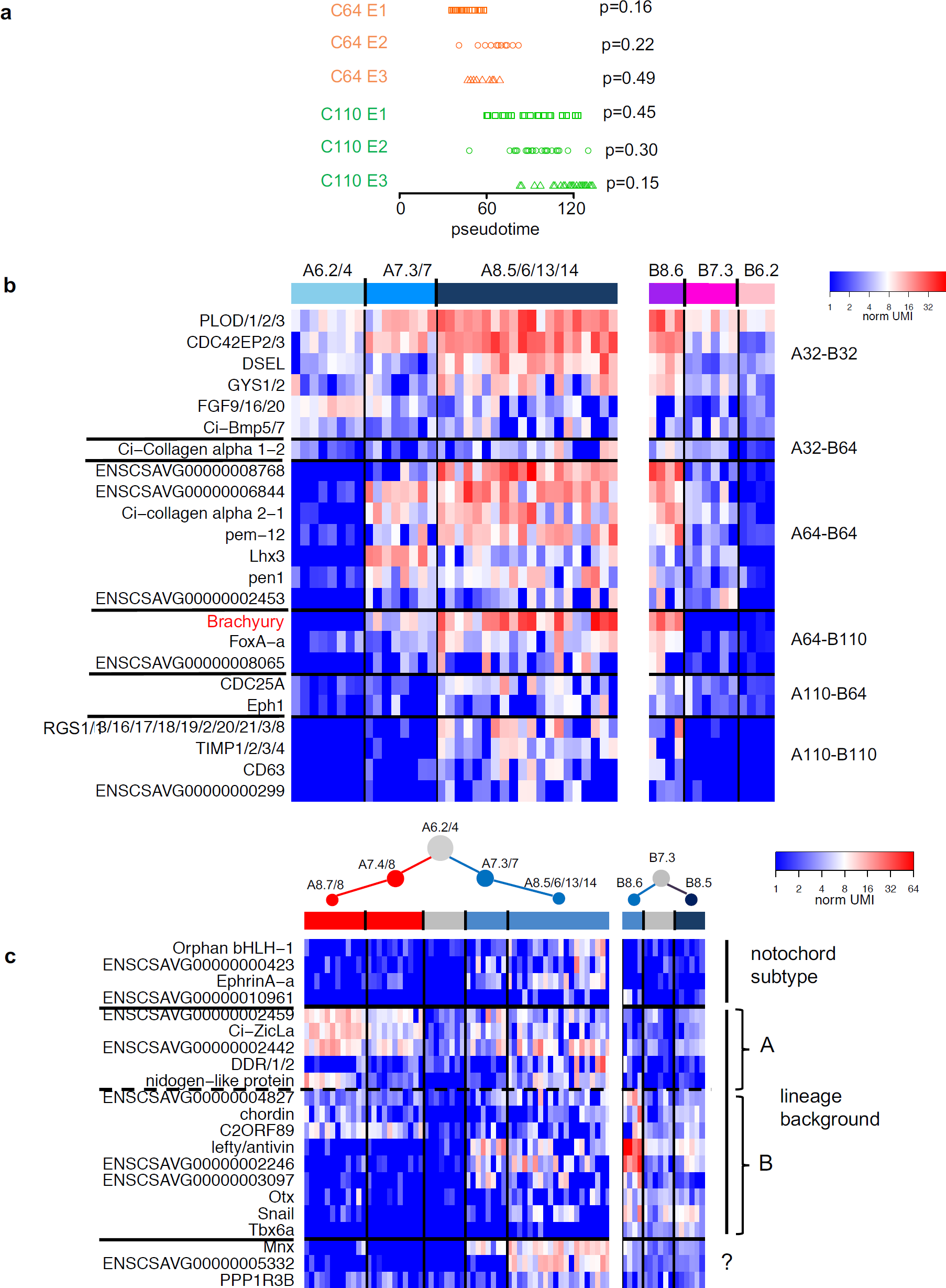
Gene expression dynamics in notochord lineage differentiation. **(a)** Order of cells along Dimension 1 in Figure 5b. Each symbol is a cell. Each embryo is shown as a row. The p-value is the probability of the orders observed in the corresponding embryo being a random ordering among 3 embryos of the same embryonic stage (C64: 64-cell stage; C110: 110-cell stage). **(b)** Expression of the 23 notochord-specific genes in the A-line and B-line notochord lineage ordered according to shared expression between A-line and B-line notochord cell stage. Heatmap follows convention in Figure 2d. Horizontal black lines separate genes of the different patterns depicted in Figure 5d. **(c)** Expression of genes that show >2 fold difference in expression levels between the A-line and B-line notochord lineage at different stages. Heatmap follows convention in Figure 2d. For the categories on the right, see main text.

**Figure S14.**
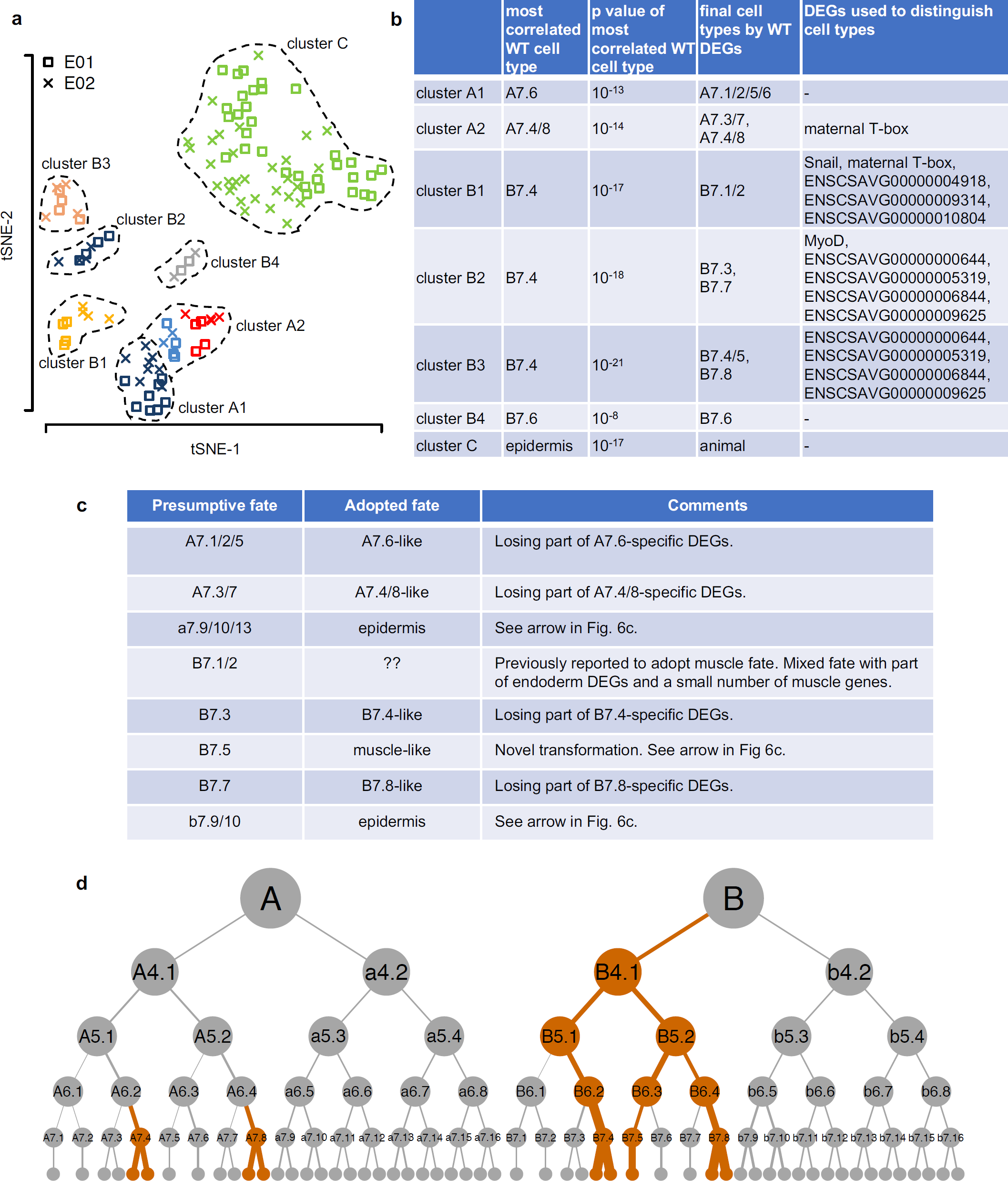
Fate transformations and asymmetry of MT genes upon U0126 treatment. **(a)** Defined cell clusters in U0126-treated embryos. Convention follows Figure 2a. **(b)** Blastomere identity assignment process for defined cell clusters. **(c)** List of detected fate transformations with complications discussed in comments and main text. **(d)** Lineage asymmetry of MT-coded genes. Cell with above background level of MT-coded genes are colored orange. Thickness of the line above an orange cell proportional to the relative expression level of MT-coded genes in the cell (fold over background of its generation). Cell names shown for 64-cell stage and earlier.

**Figure S15.**
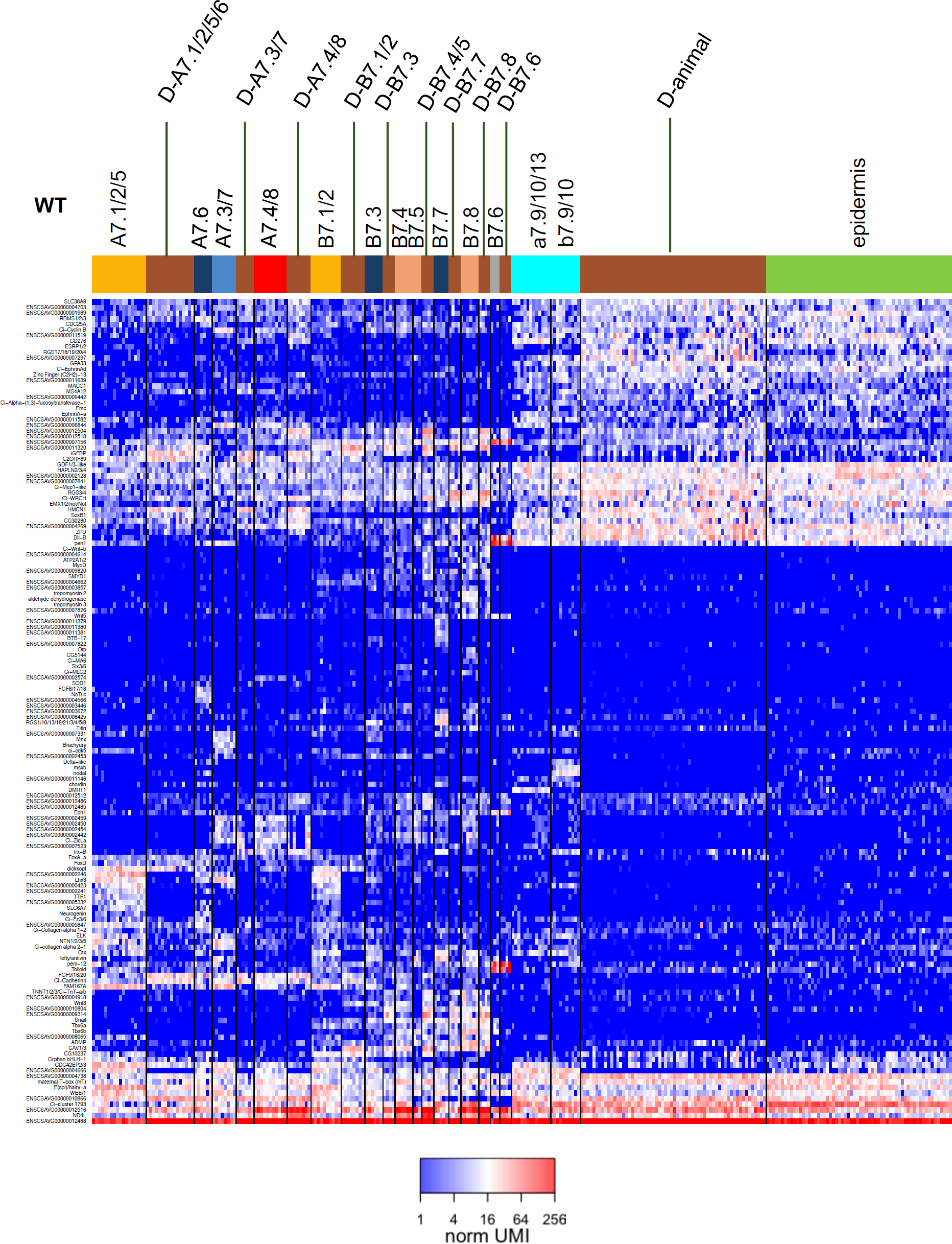
Comparison of DEG expression between wild-type and U0126-treated embryos. Heatmap convention follows Figure 2d. Genes for comparison are DEGs defined in the wild type 64-cell embryo.

